# IL-10 driven memory T cell survival and Tfh differentiation promote HIV persistence

**DOI:** 10.1101/2021.02.26.432955

**Authors:** Susan Pereira Ribeiro, Malika Aid, Frank P. Dupuy, Chi Ngai Chan, Judd Hultquist, Claire Delage, Eirini Moysi, Deanna Kulpa, Lianghzu Li, Xuan Xu, Banumathi Tamilselvan, Jeffrey Tomalka, Michael Nekorchuk, Kathleen Busman-Sahay, Rebeka Bordi, Camille Simoneau, Jean Philippe Goulet, Vincent Marconi, Jean Pierre Routy, Robert Balderas, Luca Micci, Bonnie Howell, Dan H. Barouch, Nevan Krogan, Constantinos Petrovas, Mirko Paiardini, Steven G Deeks, Jacob D. Estes, Daniel Gorman, Daria Hazuda, Rafick Pierre Sekaly

## Abstract

Mechanisms regulating HIV persistence are complex and not well understood. Increased IL-10 levels were positively associated with HIV reservoir in blood and lymph nodes (LN) of treated HIV aviremic individuals. In LNs, B cells, regulatory T cells, follicular T helper cells (Tfh), monocytes and macrophages contributed to the frequencies of IL10+ cells. Cells with HIV DNA in LNs were in close proximity to IL-10+ cells and/or had the active form of STAT3, the transcription downstream of IL-10. Gene signatures and proteins associated to cell survival, Co-inhibitory receptors expression, maintenance of memory T cells, immune metabolism and Tfh frequencies were all modulated by IL-10 and associated with HIV reservoir persistence. *In vitro,* STAT3 knockout or neutralization of IL-10, reverted all the aforementioned pathways and resulted in 10-fold decay in HIV reservoir. Collectively, these results provide strong evidence for a pivotal role of IL-10 in HIV persistence, and a potential therapeutic strategy for HIV cure.

## Introduction

HIV infection triggers a cascade of cytokine production commonly referred as cytokine storm(McMichael et al., 2010, Brockman et al., 2009). Sensing of the virus by PRRs (Pattern recognition receptors)(Jakobsen et al., 2015, Decalf et al., 2017, Hertoghs et al., 2017) and concomitant early gastrointestinal tract barrier damage ensuing bacterial translocation boosts generalized inflammation and immune activation. Importantly, the sensing of bacterial products by TLRs can trigger IL-10 production(Zevin et al., 2016), a pleiotropic anti-inflammatory cytokine(Rojas et al., 2017) that is dramatically increased within weeks of the onset of HIV and SIV infections(McMichael et al., 2010, Estes et al., 2006). IL-10 is produced by several innate immune cell subsets (monocytes, macrophages, dendritic cells) and B cells downstream of the activation of PRRs including TLRs(Kubo and Motomura, 2012). T regulatory cells (Tregs) and Type I regulatory cells (Tr1) also produce IL-10 upon TCR engagement(Shive et al., 2015). IL-10 impedes crucial APC functions by inhibiting cytokine production and preventing the upregulation of Major Histocompatibility Complex (MHC) and co-stimulatory molecules(Mittal and Roche, 2015, Rojas et al., 2017). IL-10 also acts directly on T cells by inhibiting tyrosine phosphorylation downstream of CD28 signaling in T cells(Taylor et al., 2007), decreasing their proliferation, differentiation, and cytokine production (IL-2/IFNg). Several members of the Herpes virus family including CMV, EBV and others encode for IL-10 analogs and as a result, these viruses evade the immune response by inducing peripheral tolerance and promote their own survival(Weber-Nordt et al., 1996) and persist for life in infected hosts(Herbein, 2018, Jochum et al., 2012, Avdic et al., 2011, Slobedman et al., 2009). Hence in LCMV infection, production of IL-10 correlates with poor pathogen control(Kahan and Zajac, 2019); experimental ablation of IL-10 or inhibition of IL-10 signaling restores pathogen control and reduces the severity of disease(Couper et al., 2008). Furthermore, in HIV infected individuals, IL-10 is known to inhibit CD4 and CD8 T cell proliferation and cytokine production *in vitro*; blockade of IL-10 efficiently restored these functions in both HIV and HCV infected individuals *in vitro* (Brockman et al., 2009, Clerici et al., 1994, Landay et al., 1996, Wilson and Brooks, 2011, Cacciarelli et al., 1996, Rigopoulou et al., 2005, Yang, 2009).

HIV integrates into the host genome and persists in a small pool of long-lived or proliferating memory CD4 T cells(Lee and Lichterfeld, 2016, Murray et al., 2016, Persaud et al., 2000). Low levels of viral transcription/translation and molecular mechanisms that maintain the survival of productively infected cells facilitate the persistence of the HIV reservoir in the host(Chomont et al., 2009). Latently infected cells are known to express high levels of co-inhibitory receptors, such as PD-1, LAG-3, TIGIT and CTLA4(Fromentin et al., 2016, Chomont et al., 2009, Fromentin et al., 2019, McGary et al., 2017). Follicular T helper cells (Tfh) in B cell follicles also contribute to the formation and maintenance of the HIV reservoir, as they harbor quantitatively more intact provirus(Perreau et al., 2013) than non-Tfh cells. Of note, B cells which produce IL-10 are dependent on IL-10 for their own survival(McGary et al., 2017), and are critical for the maintenance of Tfh frequencies(Kerfoot et al., 2011). *Little is known about the role that IL-10 plays in the establishment and the maintenance of HIV reservoir size. Herein, we used unbiased holistic approaches, including tissue in situ, ex vivo, and in vitro experimental methodologies to identify and mechanistically interrogate the interplay between IL-10 and HIV persistence in infected patients*.

## RESULTS

### Significantly increased levels of circulating IL-10 in treated HIV infecte individuals (aviremics) is associated with the size of latent HIV reservoir

In plasma from antiretroviral (ART) treated HIV aviremic individuals (n=24) and HIV negative healthy controls (HC, n=4) (subgroup representative of a larger cohort, **Table S1**), 23 cytokines (IFN-2a, IFN-b, IFN-g, IL-10, IL-15, IL-17A, IL-18, IL-1b, IL-2, IL-21, IL-22, IL-27, IL-29, IL-33, IL-4, IL-6, IL-7, IL-8, IL-9, TNF-a, TGF-b1, TGF-b2 and TGF-b3) were evaluated using the Meso-Scale platform (MSD). IL-10 was the only cytokine with significantly higher levels in HIV aviremic individuals when compared to HIV negative healthy individuals [Median HIV-aviremics: 455 fg/mL, fold change (FC) HIV/HC: 1.4, p<0.05] (**Fig S1a and** **Fig 1a**) that was also positively correlated with the frequencies of circulating latently infected cells as measured by HIV integrated DNA (HIV IntDNA), a well-accepted readout for HIV reservoir(Eriksson et al., 2013) (Median HIV-aviremics: log 2.85 cps/10^6^ CD4 T cells, p<0.05/r=0.44 – **Fig S1b and** **Fig 1b**). None of the other cytokines known to trigger the transcription factor (TF) STAT3(Donnelly et al., 1999), such as IL-9, IL-6, IL-21, IL-22, IL-27(Demoulin et al., 1996, Hillmer et al., 2016), were increased or associated with HIV IntDNA in this cohort (**Fig S1a** and **b**, respectively). The potential role of IL-10 in the persistence of HIV was further supported by the heightened per cell expression of the IL-10 receptor (judged by Mean Fluorescence Intensity - MFI) in different memory CD4 T cell subsets from HIV aviremic as compared to HIV negative healthy individuals (Median MFI IL-10Ra Total CD4 HIV-aviremics: 6167, FC HIV/HC: 1.2, p=0.003 - **Fig. S1c**) as measured by flow cytometry. The per cell level of IL10Ra expression in CD45RA+ cells (a cell subset that includes T stem cell memory (TSCM), a target for HIV infection(Chahroudi et al., 2015)), was also positively associated with HIV IntDNA reservoir (p<0.03/ r= 0.37 - Pearson Correlation **- Fig. S1d)**. It is important to highlight that no other clinical parameter was associated with HIV reservoir in this study (time of HIV infection: median: 7.5 years – p = 0.70, r = -0.07; age: median: 50.5 years old - p = 0.75, r = -0.06; time under ART: median: 3 years - p = 0.34, r = - 0.16; CD4 nadir: median 251 cells/mL - p = 0.31, r = -0.17; CD4 counts: median 446 cells/mL - p = 0.25, r = -0.20, CD8 counts: median 662 cells/mL - p = 0.72, r = 0.06; CD4/CD8 ratio: median: 0.58 - p = 0.17, r = -0.24; sCD14: median: 1662 ng/mL - p = 0.26, r = 0.19 – data not shown). *Our data indicate that IL-10 may play a significant role for HIV persistence in ART HIV individuals*.

**Fig 1.**
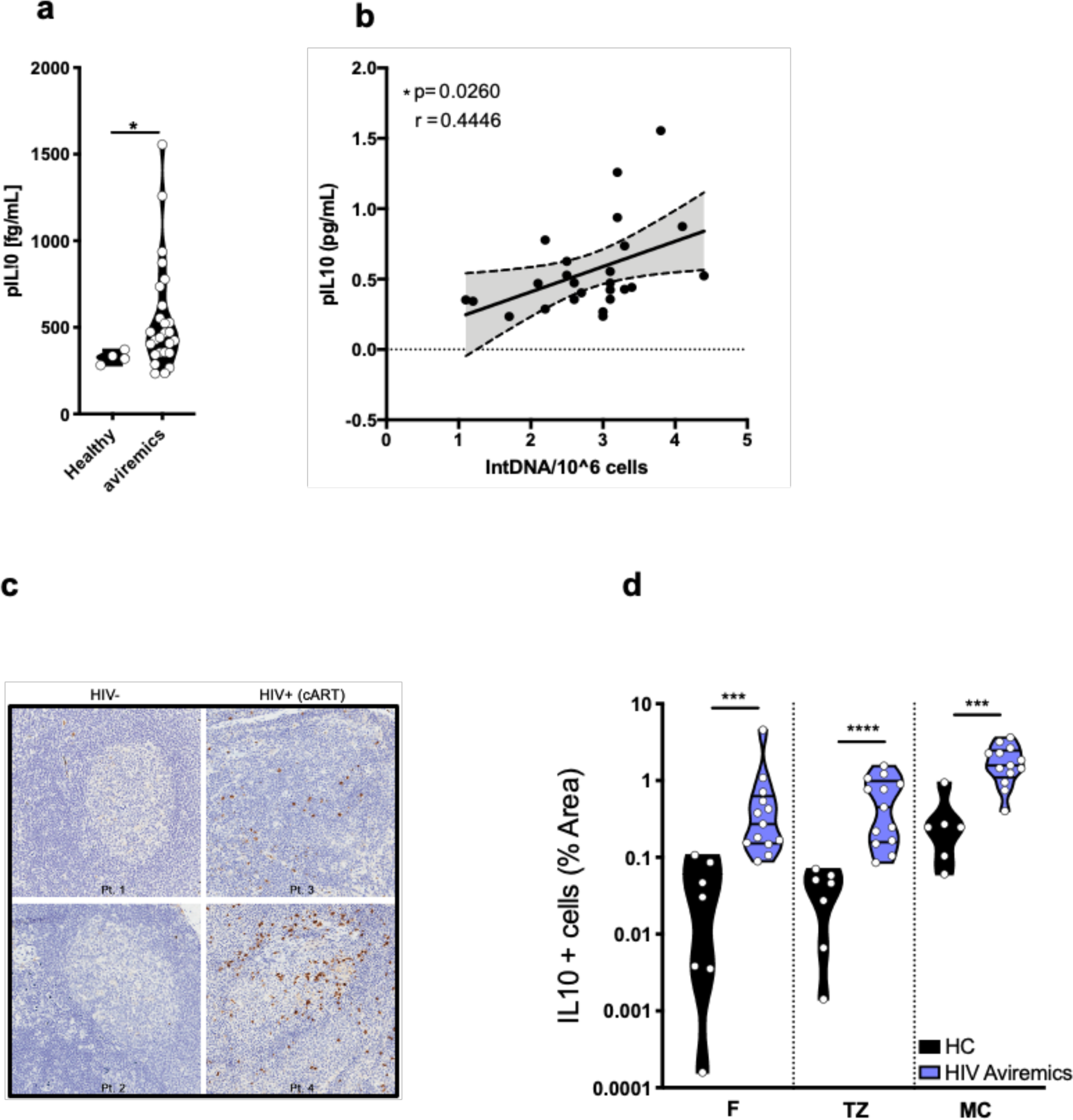
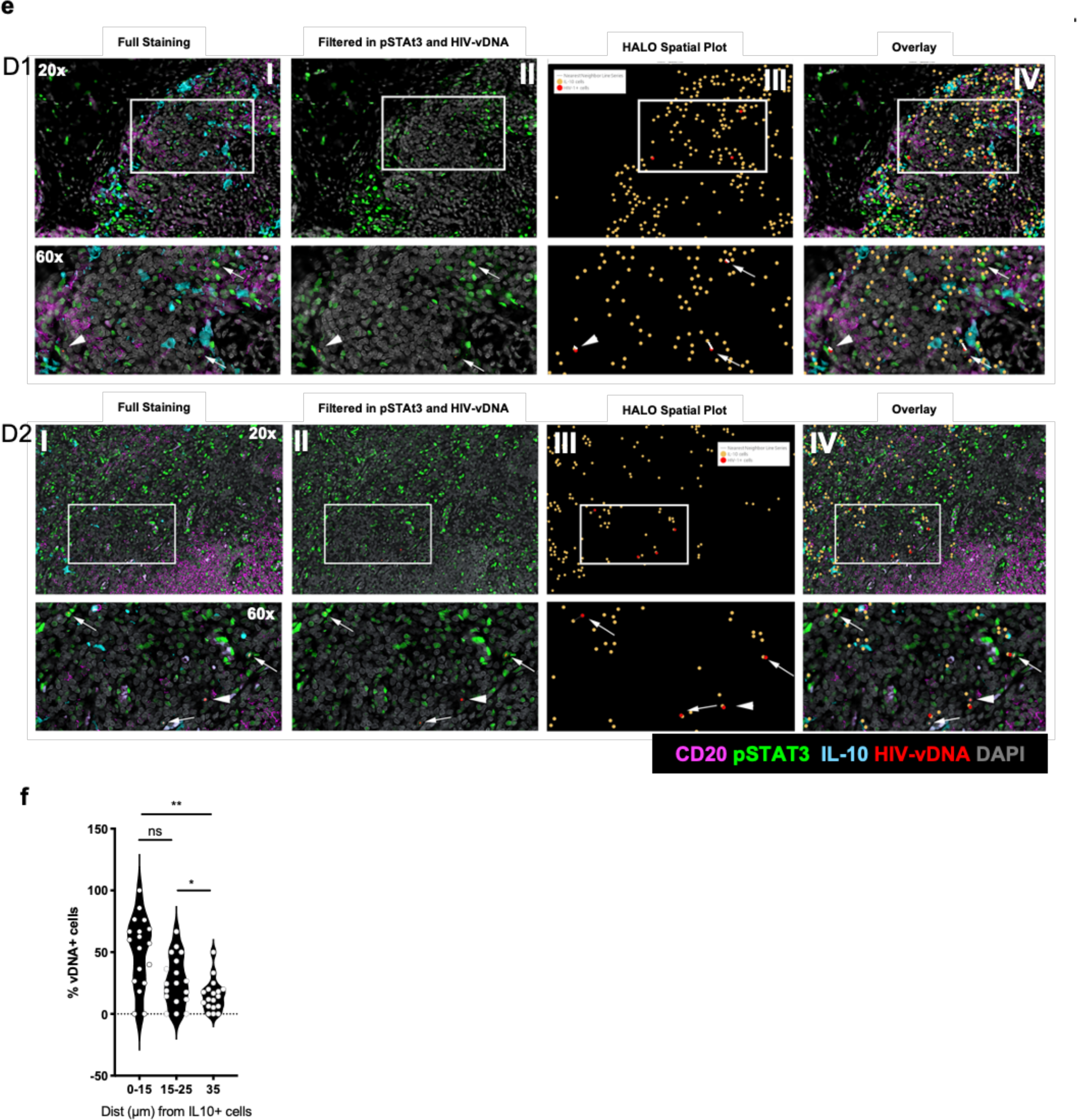
Significantly increased levels of circulating and lymph-nodes IL-10 in treated HIV infected individuals (aviremic) is associated with the size of latent HIV reservoir (HIV IntDNA). **a)** Cytokine array was performed using Meso-Scale platform (MSD). pIL-10 levels were increased in HIV-aviremic individuals (n=24) when compared to HIV negative Healthy controls (HCs, n=4); **b)** pIL10 levels were positively correlated to latent HIV reservoir as measured by HIV IntDNA; **c)** Representative image of IL-10 expression in LNs from 2 HIV negative HCs (left plots) or 2 HIV-ART individuals (right plots). Brown chromogenic IHC – IL-10+ cells –Brown; tissue color: hematoxylin staining; **d)** Quantification of IL-10+ cells (frequency of total area) in different areas of the LN in HIV negative HC (back, n=7) and HIV-aviremics (blue, n=13). F (Follicle), TZ (T cell zone) and MC (medullary cord) are shown. Mann-Whitney unpaired T test was used to compare the frequencies of IL-10+ cells between different zones: F, TZ and MC; **e)** Representative DNAscope (vDNA: red), followed by pSTAT3 (green), IL-10 (cyan), CD20 (pink) and DAPI (grey) multiplexed immunofluorescence in two HIV- aviremic. For each subject (1 and 2) the top row (magnification of 20x) 4 plots represent: I - Full staining (overlap of all markers), II – Filtered in pSTAT3 and HIV-vDNA, III – HALO spatial plot, IV – Overlay of I and III. Arrows point to vDNA+P-STAT3+ and arrowheads point to vDNA+P-STAT3-cells. Nearest neighbor and Proximity detection of HIV-1 vDNA+ cells to nearby IL-10+ cells was calculated (plot III). The Spatial Proximity Map (III) shows the location of IL-10+ cells (Yellow dots) and HIV-1 vDNA+ cells (Red dots) as detected and assigned by the HALO software. The nearest IL-10+ cell to each HIV-1 vDNA+ cell is indicated by the Proximity Line (gray); **f)** Frequencies of HIV DNA+ cells in the nearest IL10+ cell area is calculated based on spatial proximity map (III). NS: not significant, *p<0.05, **p<0.01. LN: lymph nodes; IHC: Immunohistochemistry; FC: Fold Change. 5-7uM: diameter of a single cell.

### Significantly higher HIV DNA+ cells in lymph nodes from HIV aviremic individuals are associated with close proximity to IL-10+ cells

We next used several *in situ* quantitative imaging approaches to evaluate the contribution of IL-10 to the cellular and anatomical localization of the HIV reservoir in lymph nodes (LNs). In LNs from HIV negative HC (left plots, n=7) and HIV-aviremics (right plots, n=13), *in situ* immuno-histochemistry (**Fig 1c**) revealed that the frequencies of IL-10+ cells per anatomical area, *i.e* the Follicle (F), T cell zone (TZ) and medullary cord (MC) were significantly increased in HIV aviremic individuals when compared to HC (F: *HIV aviremics: 0.27% IL-10+ cells, FC HIV/HC: 9, p<0.0001*, *TZ: HIV aviremics: 0.46% IL-10+ cells, FC HIV/HC: 9.8, p<0.0001; MC: HIV aviremics: 1.6% IL-10+ cells, FC HIV/HC: 6.4, p=0.0003* – **Fig. 1d** – Mann-Whitney unpaired T test). The contribution of the IL-10 pathway engagement in the maintenance of the HIV reservoir in follicular (F) and extra-follicular (EF) areas of LNs was further evidenced by the proximity of IL-10+ cells to HIV vDNA+ cells, as well as the proportion of vDNA+ cells that expressed phosphorylated STAT3+ (pSTAT3) cells. DNAscope (vDNA: red), followed by pSTAT3 (green), IL-10 (cyan), CD20 (pink) multiplexed immunofluorescence staining was performed. Representative images from 2 donors show the expression of the selected markers in the follicle (top panel– donor 1) and in the extra-follicular area (bottom panel – donor 2) (**Fig 1e**). While the overall frequency of pSTAT3+ cells was on average 11.3% (SEM ±1.8) in all LN compartments [8.8% (SEM ±1.3; F), 10.6% (SEM ±1.7; EF) and 14.5% (SEM ±2.1; MC)], the frequency of HIV vDNA+ cells that were pSTAT3+ was on average 30.1% (SEM ±5.1) in the LNs of HIV aviremic individuals (n=18) (average quantification from all images as in **Fig 1e** for all donors evaluated). *Importantly, the frequencies of HIV vDNA+ cells increased significantly with their proximity to IL-10+ cells* (0-15uM Median: 60% HIV vDNA+, 15-25uM Median: 25% HIV vDNA+, 35uM Median: 13% HIV vDNA+ cells; 0-15um vs 15-25uM p>0.05; 15- 25uM vs 35uM p<0.05, 0-15um vs 35uM p<0.01 – **Fig 1f**) as measured by HALO (Spatial Analysis plot - detailed in methodology). Importantly, 92.6% of all HIV vDNA+ cells were within an average distance of ∼3-5 cell diameters (35um) from an IL-10+ cell (quantification from **Fig 1e -** n=18 donors). Our results capture a static snapshot of the LN structure; hence these findings most likely represent an underestimation of the number of infected or IL-10+ cells, as cells in lymph nodes are in constant movement(Huang et al., 2004, Bajenoff et al., 2007, Germain et al., 2012). *Nevertheless, these data strongly and significantly infer the importance of IL-10 signaling in HIV persistence in LNs*.

### Several lymphoid and non-lymphoid cell subsets contribute to IL-10 production in LNs of treated chronic aviremic HIV-infected individuals

Using multiparametric confocal imaging and histo-cytometry(Gerner et al., 2012) we demonstrate that several cell subsets including monocytes and macrophages **(Fig S2a** and **Fig 2a**) as well as CD4 T cell subsets and B cells (**Fig S2b** and **Fig 2b**) contributed to IL-10 production in the F and EF areas of LNs. A representative position map is shown in **Fig 2c**. The EF zone presented higher frequencies of IL-10hi cells than F zone (Images in **Fig 2a-b** – IL-10 red dots, quantification in **Fig 2d**-p<0.05, FC EF/F: 4.05, n=5). Considering the relative frequencies of IL10hi cells in the F and EF areas in our cohort, IL-10 expressing cells included CD163+/CD68, CD163-/CD68+ and CD163+/CD68+ monocyte/macrophage; the latter cell subset contributed to significant higher frequencies of IL-10 + cells in the EF area than in Follicles (*p<0.0001, 1.74-FC increase into EF compared to follicular area*-**Fig 2a and e, n=5**, gating strategy **Fig S2a).** Although most of IL-10+CD4 T cells were FoxP3-, a readily fraction was found to be FoxP3+ in the F and EF areas (*Foxp3+: FC EF/F: 5.72, p=0.59; Foxp3-: FC EF/F: 1.37, p= 0.45* (**Fig 2b, e**, n=4, gating strategy **Fig S2b)**. In the B cell follicle (BCF), regular immunohistochemistry (IHC) multiplexing markers for B cells (CD20: blue), CD4 cells (red), PD-1 (orange) and IL-10 (green), we observed that Tfh cells (CD4+ PD1+(white circles, **Fig 2f**), contributed to 42% of the IL10+ cells in the BCF (**Fig 1g,** n=11). *Thus, several LN cell subsets can contribute to IL-10 production and could impact on HIV reservoir persistence in this relevant compartment in vivo*.

**Fig 2.**
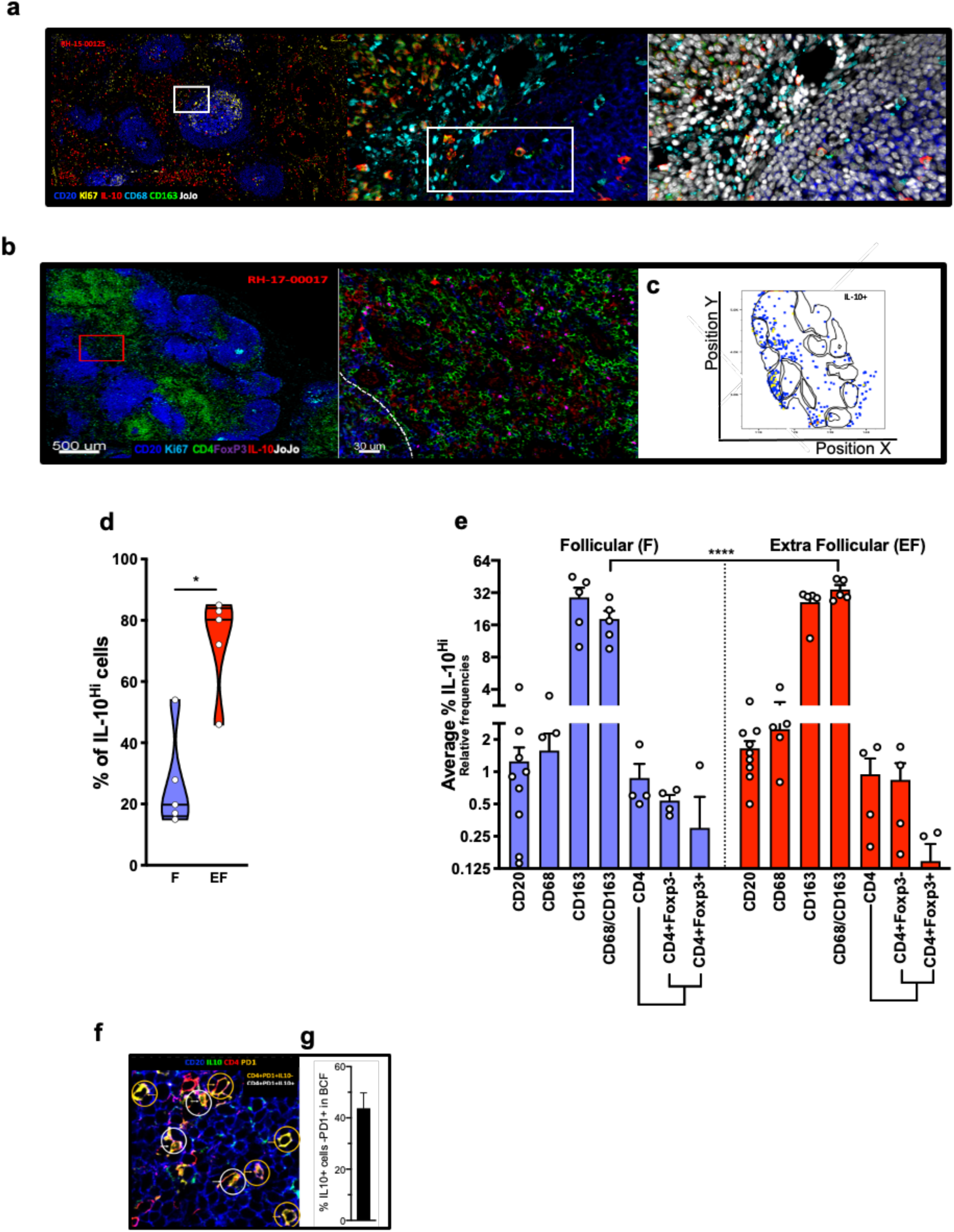
Several lymphoid and non-lymphoid cell subsets contribute to IL-10 production in LNs of treated chronic aviremic HIV-infected individuals: Monocytes, macrophages, CD4 T cells (Tregs, non-Tregs), B cells and Tfh cells are IL10+ in the LNs from HIV-aviremics individuals. **a)** Representative confocal images (40x) of an LN section of a treated aviremic HIV+ individual showing staining for CD20 (blue), Ki67 (yellow), IL-10 (red), CD68 (Cyan), CD163 (green) and the nuclear stain JoJo (upper row and left panel). Zoomed-in details (upper row - middle and right panels) and single-color captions (bottom row - right panel) are from a representative example of CD68/CD163/IL-10 staining that was used for the quantitation. Zoomed-in areas are presented in succession and each zoom-in corresponds to the region demarcated by the white enclosure. **b)** Representative confocal images (40x) of LN sections of two treated aviremic HIV+ individuals. Overviews on the left depict the total LN area imaged and distribution of CD20 (blue), Ki67 (cyan), CD4 (green) in the tissues screened. Zoomed-in details show the distribution of CD20 and CD4 in overlay with IL-10 (red) and FoxP3 (magenda) at the follicular (CD20^hi/dim^) and extrafollicular (CD20-) junction as denoted by the white dotted lines. Zoomed areas correspond to the red squares shown in each tissue overview; **c)** IL10 location plot; **d)** Pooled histo-cytometry data (n=5) showing the average frequency of IL-10^hi^ cells in follicular (CD20^hi/dim^: average 26.75%) versus extra-follicular (CD20^-^: 73.25%) areas as assessed by multiparameter confocal imaging (FC EF/F: 3, p<0.05); Gating strategy in **Fig S2**; **e)** Contribution of each of the subsets evaluated by confocal/histo-cytometry for the total frequencies of IL10+ cells in the F and EF. CD20+, CD68+ CD168+, CD163+CD68+, total CD4, CD4+Foxp3-, CD4+Foxp3-, co-stained for IL-10 are shown contributing to 100% cells IL10+ in each area; **f)** Representative IHC in BCF from one HIV aviremic individual: CD20 (blue), IL10 (green), CD4 (red) and PD1 (Orange) staining. White and orange circles highlight CD4+PD1+IL10+, CD4+PD1+IL10-cells, respectively; **g)** Frequencies of CD4+PD1+IL10+ Tfh cells (white circles) are shown.

### Pathways associated with cell survival, T cell memory maintenance, co-inhibitory receptors expression, metabolism and Tfh cell differentiation are positively associated with pIL-10 levels and HIV IntDNA

Transcriptional profiling was used to identify the mechanisms downstream of IL-10 that underlie its association with HIV persistence. Gene expression profiles of whole blood from HIV-aviremic individuals were correlated by linear regression analysis with the frequencies of cells harboring HIV IntDNA and to pIL-10 levels (**Fig 3a,** n=22). Gene Set Enrichment Analysis (GSEA)(Subramanian et al., 2005) showed that pathways downstream of IL-10/STAT3 signaling were positively correlated to pIL-10 (NES: 2.16, FDR<10-6, p<10-6) and HIV IntDNA levels (NES1.79, FDR<10-6, p<10-6) (**Fig 3b**; Leading edge genes - LEGs, **Fig S3a)**. Additionally, correlated pathways and genes encoded for molecules with anti-apoptotic properties which are all targets of the JAK-STAT signaling pathway (BIRC5 FYN, BCL2L1, NOTCH, MYC; pIL-10 NES: 1.9, FDR<10-6, p<10-6; HIV IntDNA NES: 1.88, FDR<10-6, p<10-6), as well genes endowed with antiapoptotic functions (BCL2L1, PIM1) or that act as regulators of cell metabolism (MYC, JUN)(Wang et al., 2011), genes related to glycolysis and glucose catabolic process (STIP1, ENO1, GAPDH - pIL- 10 NES: 2.15, FDR<10-6, p<10-6; HIV IntDNA NES: 1.99, FDR<10-6, p<10-6), genes associated to the maintenance of Central Memory T cells, the long-term reservoir (TCM) (CXCR3, EHD4, TOP2A, TNFRSF4 - pIL-10 NES: 3.44, FDR<10-6, p<10-6; HIV IntDNA NES: 2.37, FDR<10-6, p<10-6), genes encoding for co- inhibitory-receptors (LAG3, TIGIT, PDCD1, CTLA4; pIL-10 NES: 2.04, FDR<10-6, p<10-6; HIV IntDNA NES: 2.03, FDR<10-6, p<10-6) and their downstream signaling targets (PIM3, CASP3, PRMT1, NFKBIZ, KAT2A), as well as genes that define Tfh differentiation (BCL6, MAF, CXCR5, ICOS-ICOSL pIL-10 NES: 1.7, FDR<10-6, p<10-6; HIV IntDNA NES: 1.39, FDR= 0.005, p<10-6) (**Fig 3b** **–**pathways; and **Fig S3a** – LEGs) were all associated to pIL10 and HIV IntDNA levels (**Table S2 -3**). *These results highlight the contribution of several biological processes (cell survival, induction of Co-IR, cell, memory maintenance, metabolism and Tfh differentiation), importantly STAT3/IL-10 signaling, as associated to ex vivo plasma levels of IL-10, as potential drivers of HIV reservoir persistence*.

**Fig 3.**
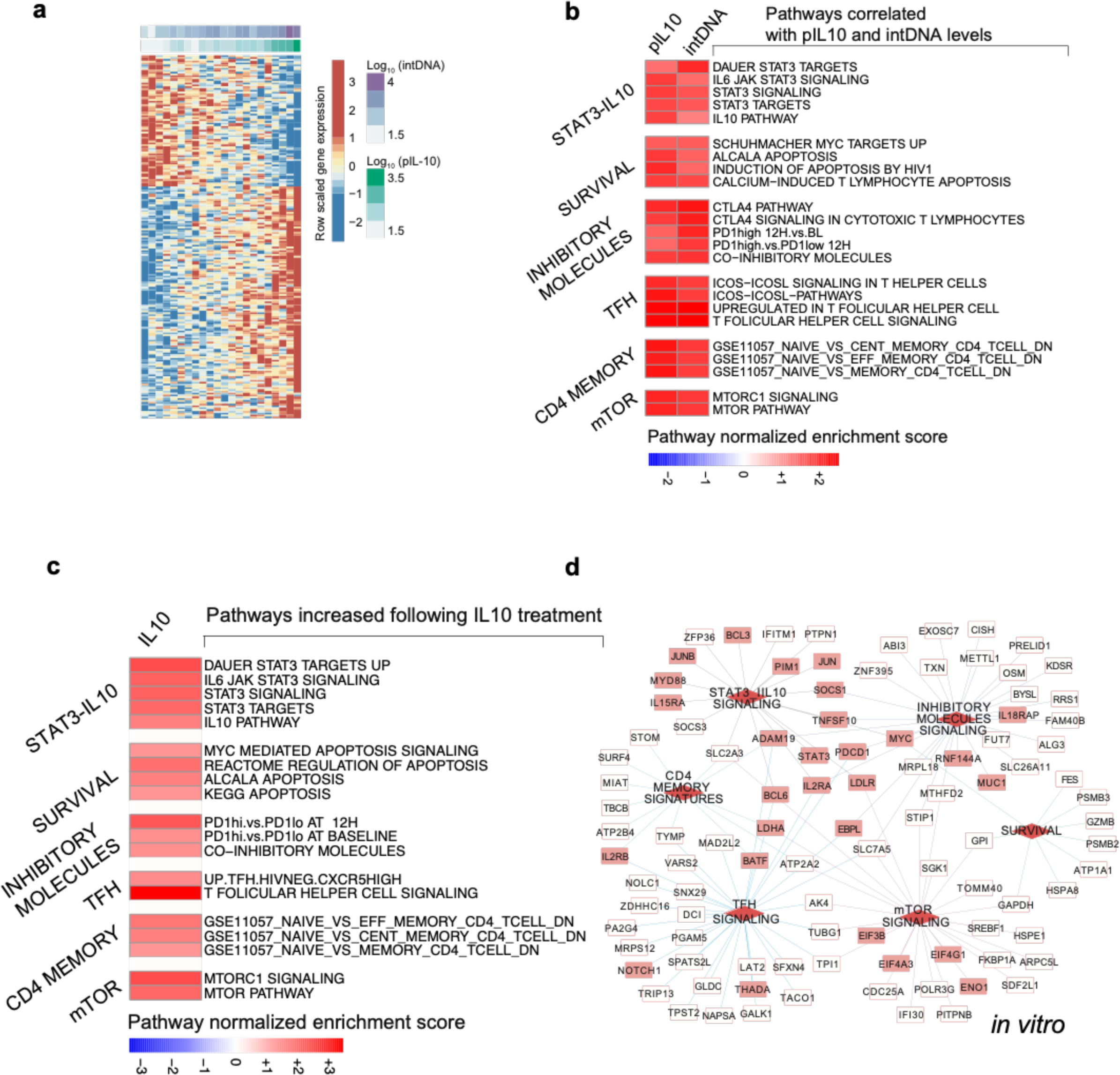
Pathways associated with cell survival, T cell memory maintenance, co-inhibitory receptors expression, metabolism and Tfh cell differentiation are positively associated with pIL-10 levels and HIV IntDNA and specifically induced by IL-10 stimulation in vitro. Gene array was performed in whole blood from HIV aviremic individuals. **a**) Linear regression model was performed for each gene with pIL-10 or HIV intDNA as an independent variable. Gene expression as a dependent variable was fit using the R package LIMMA. Genes that correlated with both outcomes using a Benjamini– Hochberg corrected p value of 0.05 were selected. The top 100 genes were plot using pheatmap R package. Rows represent genes and columns represent samples. pIL-10 and HIV IntDNA were used as continuous variables and their log 10 transformed values were shown on the top of the heatmap; **b)** Gene sets enrichment analysis (GSEA) in *ex vivo* samples revealed significant and positive correlation of gene signature with HIV IntDNA and pIL-10 levels. Gene signatures were compiled from MSigDB C2 and C7 and other gene sets available in the literature. The LIMMA package was used to fit a linear regression model to each probe (log2 expression) to respectively HIV IntDNA and pIL-10 levels. Genes that correlated to the outcomes were sorted from high to low using their coefficient correlation for each regression and then submitted to GSEA using the pre Ranked list option. Signatures significantly enriched in these lists with an FDR cut-off of 5% were selected. GSEA normalized enrichment scores were plotted using pheatmap R package. Color gradient from blue to red depicts the normalized enrichment score (NES) ranging from decreased (blue) to increased (red); **c)** IL10 stimulation *in vitro* induces the same pathways associated with higher levels of HIV IntDNA and pIL-10 in chronic HIV infected individuals. Gene array was performed in CD4+ T cells isolated from healthy donors and stimulated for 12 hours with IL-10 or left unstimulated. Gene induced by IL-10 stimulation *in vitro* at 12 hours compared to unstimulated cells were identified using the LIMMA R package and an adjusted p value cut-off of 0.05. GSEA as described above was used to identify which gene sets are enriched in IL-10 stimulated cells. Gene sets NES of pathways significantly induced by IL-10 are shown in the heatmap. Color gradient from blue to red depicts the enrichment score ranging from decreased (blue) to increased (red). For both panels b and c, pathways involved in the same signaling were grouped into modules of STAT3_IL10 signaling, survival, Tfh signaling, inhibitory molecules signaling, memory signatures and metabolism; **d)** Network representation of Leading-edge genes (LEGs) that contributed significantly to the enrichment of pathways induced by IL10 and shown in panel **c**. Dynet application implemented under Cytoscape was used to generate the network. Red diamonds nodes represent modules name. The edge colors connect genes from each module (STAT3/IL10 signaling: black; Inhibitory molecules signaling: dark blue; CD4 memory signatures: light blue; Tfh signaling: indigo blue; mTOR: purple; Survival: green). Rectangles represent the genes upregulated by IL-10. Highlighted as full red rectangles represent important functional gene for each module.

### IL-10 stimulation in vitro triggers the upregulation of the pathways associated with heightened pIL-10 levels and to the magnitude of the HIV reservoir ex vivo

To confirm the specific role of IL-10 in the induction of the aforementioned pathways associated with HIV reservoir persistence *in vivo*, gene array analysis followed by GSEA was performed on *in vitro* IL-10 stimulated CD4+ T cells from healthy donors. IL-10 induced pathways (**Fig 3c**) and LEGs (**Fig 3d) (Table S4)** were the same as the ones significantly correlated to *ex vivo* levels of pIL-10 and HIV IntDNA. They included the IL-10-transcription factor STAT3, its target genes as well as genes downstream of IL-10 signaling pathway (PDCD1, SOCS3, BCL3, MYC, IFITM1, BCL6,JUN, SOD2, STAT3) (NES: 2.72, FDR<10- 6, p<10-6), genes encoding for survival and anti-apoptotic pathways (PIM1, NOTCH, MYC, BCL2L - NES: 1.99 FDR<10-6, p<10-6), maintenance of T cell memory ( PIK3CA, IL7R, KLRB1, BTG1 - NES: 1.62 FDR= 6x10-4 p= 0.005), PD1 signaling pathway and exhaustion (PDCD1, CTLA-4, CASP3, BTF3, KAT2A), (PRDX4, ELMO1, IRF4) - NES: 1.69 FDR=5 x 10-4, p<10-6), maintenance of cell metabolism (ALDOA, ENO1, PKM2, LDHA, SGK1,GAPDH, PRDX1 - NES: 1.93 FDR=3.3 x 10-4, p<10-6), and Tfh signaling (BATF, BCL6, ICOS, - NES: 2.31 FDR< 10-6, p<10-6). Importantly, we found that 453 out of 644 IL-10 induced genes that correlated with the frequencies of cells with HIV IntDNA were also transcriptionally regulated by STAT3, *which highlights the restrict regulation of these pathways by IL10/STAT3 signaling.* These STAT3 target genes (ref: (http://www.beaconlab.it/ pscan) also covered all the aforementioned pathways listed as associated with pIL-10 and HIV IntDNA *in vivo* (**Fig S3b** and **c**, respectively – NES and FDR indicated in the figure).

### Soluble CD14 (sCD14) is increased in HIV infected individuals and is associated with heightened pIL-10 levels and major pathways associated with HIV persistence in vivo

Bacterial translocation occurs early upon HIV infection(Marchetti et al., 2013) as a consequence of the loss of Th17 cells in the gut(Bixler and Mattapallil, 2013). sCD14 is a validated biomarker for bacterial translocation(Kelesidis et al., 2012) as it is released from activated monocytes upon engagement to lipopolysaccharide (LPS). The engagement of bacterial Pattern recognition receptor (PRRs) to Pathogen-associated molecular pattern (PAMPs) is known to trigger IL-10 production(Yanagawa and Onoe, 2007). Indeed, in our HIV cohort of HIV-aviremics individuals, the plasma levels of sCD14 were significantly increased when compared to HIV negative HCs (HC, n=10, HIV, n=42- FC HIV/HC: 1.20, p<0.05 – **Fig S4a**) and were significantly associated with pIL- 10 levels (p=0.007/r=0.53 – **Fig S4b**) indicating that sCD14 could constitute one of the upstream signals leading to heightened pIL-10 levels in these individuals. Supporting this hypothesis, all the pathways associated with heightened pIL-10 and HIV reservoir *in vivo*, were also significantly associated with sCD14 levels (IL- 10/STAT3 signaling (OASL, IL-10RA, IL4, JAK2 - NES: 2.34, FDR<10-6, p<10-6), survival and anti-apoptosis (TNFR1B, MYD88, GAPD4 - NES: 2.05, FDR<10-6, p<10-6), maintenance of memory T cells (IL10Ra, TIGIT, TNFRb, SMAD3 - NES: 3.49, FDR<10-6, p<10-6), inhibitory molecules signaling (CTLA4, TIGIT, FYN, BHLHE40 - NES: 2.29, FDR<10-6, p<10-6), maintenance of cell metabolism (ENO1, GAPDH, SLC7A5 - NES: 2.54, FDR<10-6, p<10-6), and Tfh signaling (CXCR5, CD2, PRDM1 - NES: 1.93 FDR< 10-6, p<10-6)) (Pathways and LEGs - **Fig S4c-d** respectively, **Table S5**). Of note, several transcription factors (TFs) known to bind directly or indirectly to the IL-10 gene promoter(Kubo and Motomura, 2012, Zhang and Kuchroo, 2019, Tsuji-Takayama et al., 2008, Cao et al., 2005) including STAT3, BCL-6, MAF, MYC were among the top genes that contributed to the enrichment of these pathways and were significantly associated with sCD14 and pIL-10 levels (**Fig S4e**). Overall, 20 pathways were commonly associated with HIV IntDNA, pIL-10, sCD14 and specifically induced by IL-10 stimulation in vitro (**Fig S4f, Table S6**). These pathways included all the aforementioned biological processes (survival, co-IR, cell memory, metabolism and Tfh differentiation). *These data indicate that sCD14 could be one of the mechanisms upstream of IL- 10 production in HIV-aviremic individuals, contributing to the modulation of pathways associated with HIV persistence*.

### STAT3 knockout (KO) leads to decreased viral reservoir

To validate experimentally the mechanisms identified by transcriptional profiling associated with HIV persistence and specific to IL-10 signaling we used an *in vitro* model of HIV infection and latency (LARA: Latency and Reversion Assay(Kulpa et al., 2019)) (**Fig S5a**). In this assay, isolated memory CD4+ T cells from healthy individuals are infected *in vitro and* induced to latency by adding TGFb which our group previously established as an inducer of HIV latency *in vitro(Kulpa et al., 2019)*. To confirm the role of IL-10 in the induction of latency, we initially compared it to TGFb. As readout we have evaluated HIV protein expression (p24 – HIV gag) by flow cytometry (**Fig S6a, Table S7**) at different time-points. Addition of IL-10 after *in vitro* infection led to a significantly faster HIV p24 protein expression decay when compared to TGFb [Day 12, Fold Decay (FD) (FD IL-10/TGFb):2.21, p<0.05, Day 15, FD1.54, p<0.05, Day 17, FD2.85, p<0.05 - **Fig 4a***]. Importantly, frequencies of cells with latent HIV provirus, as measured by HIV IntDNA (n=5), were comparable between the two tested conditions (p>0.05 -* ***Fig 4b****), validating IL-10 as an inducer of latency*.

**Fig 4.**
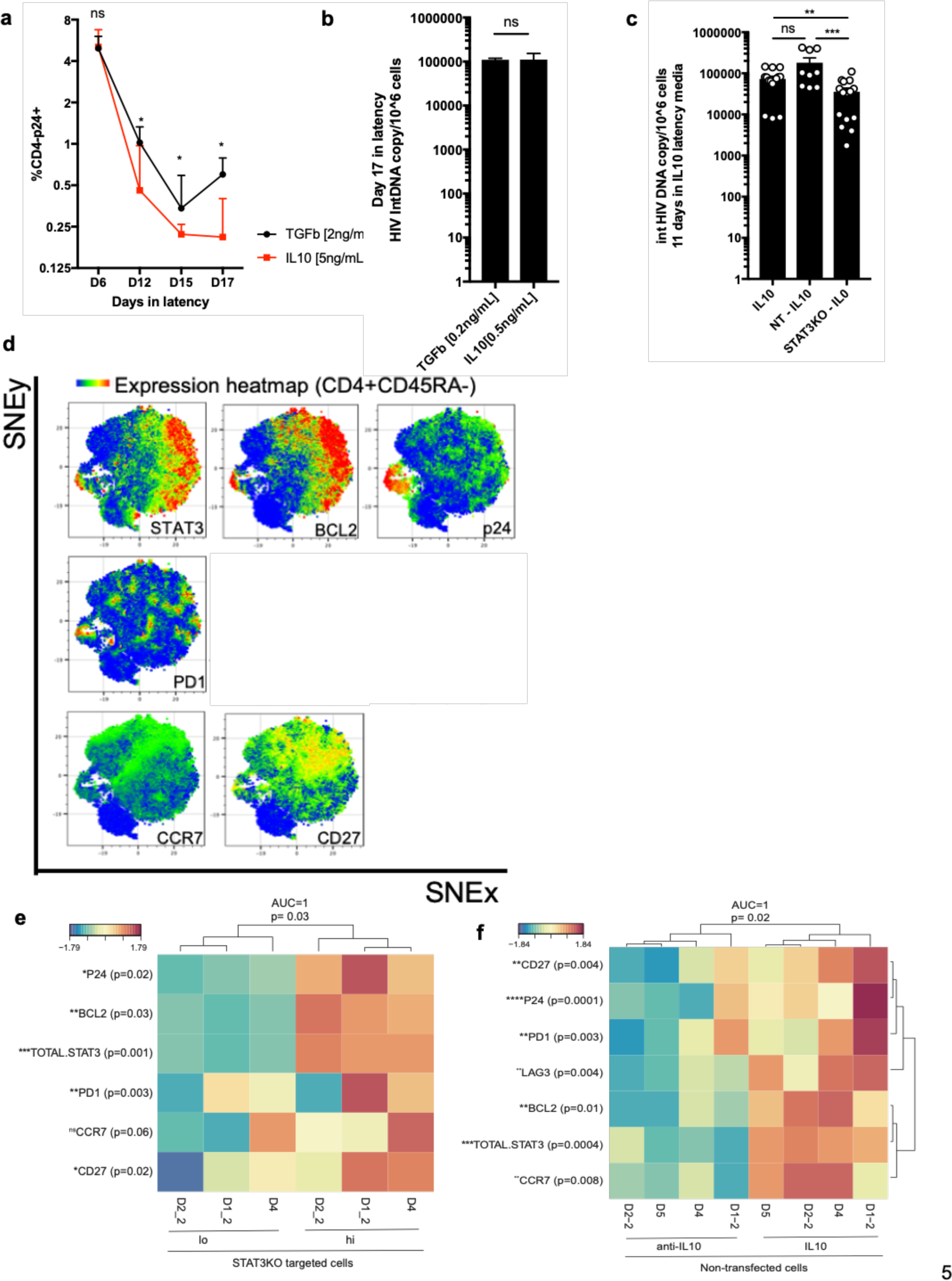
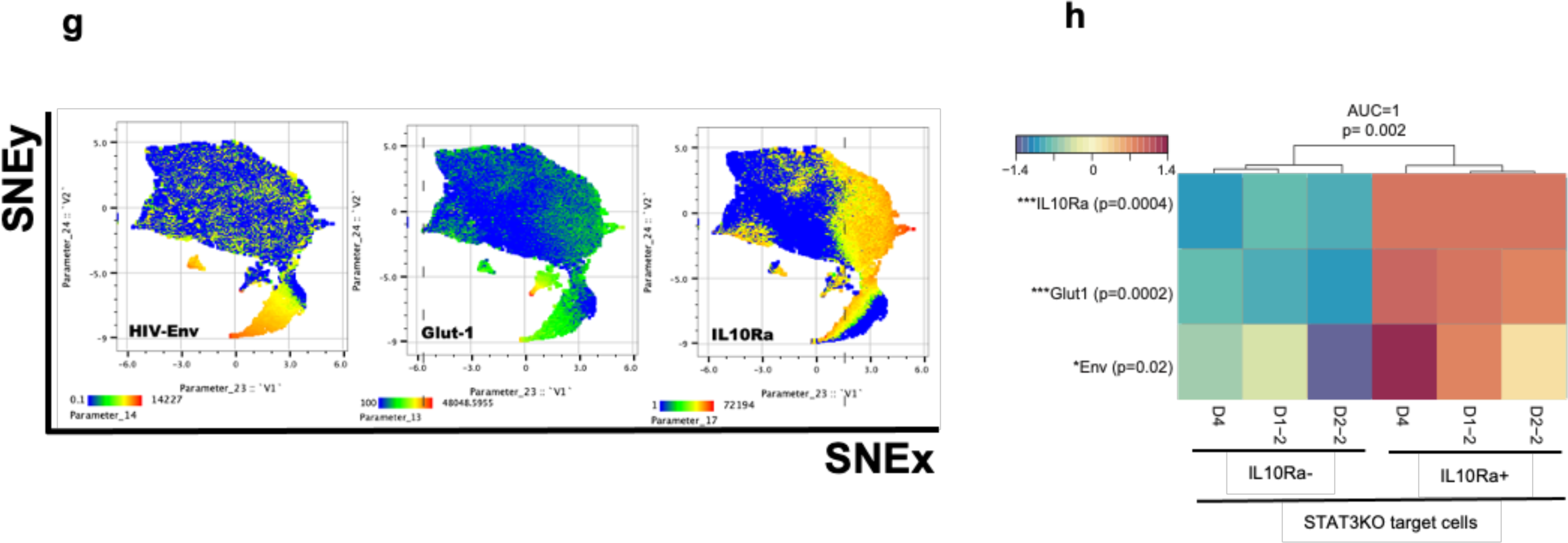
STAT3 knockout (KO) leads to decreased viral reservoir and to decreased expression of major proteins of key pathways associated with HIV persistence in vivo. Memory CD4+ T cells were isolated from HIV negative healthy controls, infected as in **Fig S3a**, and latency was induced at D3 using IL- 10 (5ng/mL) or TGFb1 (20ng/mL) in media containing H80 cell supernatants(Kulpa et al., 2019), ARVs and IL-7 (40ng/mL). **a)** Frequencies of HIV infected cells (CD4- p24+) were evaluated at (day:D) D6, D12, D15 and D17. IL-10 induces HIV protein expression decay at significantly higher levels than TGFb; **b)** HIV IntDNA copies per/million CD4 T cells evaluation at D17 (ns: non-significant); **c)** Copies of HIV IntDNA per million CD4 T cells after latency induction comparing all conditions cultured in media containing IL-10 (10ng/mL) + IL7 (40ng/mL and ARVs): non- transfected, NT and STAT3KO. Paired t test between conditions: **p<0.01, ***p<0.001. **d)** Expression of major proteins of each of the pathways associated *ex vivo* with pIL-10 and HIV IntDNA levels were evaluated by flow cytometry in CD4 T cells from LARA as described in **Fig S3a** (STAT3, BCL2, HIV-p24, PD1, CCR7 and CD27). TSNE analysis was performed in all samples in a total of 3000 cells/sample CD45RA- gated as in **Fig S4b**. TSNE-heatmap for each marker is shown. Low to high levels of protein expression are depicted in the gradient from blue (low) to red (high). **e)** Heatmap summarizing the findings in the STAT3KO transfected condition comparing STAT3hi to STAT3lo. Each row refers to a specific marker and the p values associated to each of them based on paired t. test comparing STAT3lo x STAT3hi. **f)** Heatmap summarizing the findings in the non- transfected condition comparing IL10 and anti-IL10 containing media conditions. Each row refers to a specific marker and the p values associated to each of them based on paired t. test comparing IL10 x anti-IL10. **g)** Expression of metabolic markers along with HIV-envelope expression was assessed in a second flow panel. Same analysis strategy was applied as in **d).** UMAP-heatmaps show the gradient of expression oh each of the proteins independently. **h)** Heatmap summarizing the findings in the IL10R+ and IL10R- from STAT3KO transfected condition. Each row refers to a specific marker and the p values associated to each of them based on paired t. test comparing IL10R+ and IL10R-. *p<0.05, **p<0.01, ***p<0.0001, ****p<0.0001. Gate on IL10Ra expression was used since no intracellular staining for STAT3 was performed in this panel.

The importance of the IL-10/IL-10Ra-STAT3 axis for the maintenance of HIV (p24) and for the expression of major proteins of each of the pathways associated with HIV IntDNA *ex vivo* that were as well specifically induced by IL-10 *in vitro*, was validated by the blockade of the IL-10 pathway. For this purpose, we either knocked-out STAT3 by CRISPR-CAS9 editing (STAT3 KO)(Hultquist et al., 2019) or neutralized IL-10 by using an anti-IL-10 monoclonal antibody (mAb)(L., 2005). Addition of anti-IL-10 mAb led to a significant decrease in the phosphorylation of STAT3 even at concentrations as low as 0.1ug/mL (p<0.05 - **Fig S5b**). Efficiency of STAT3 knock-out was validated in aCD3/CD28 pre-activated CD4 T cells. The transfection of pre-activated cells led to significant knockout of STAT3 protein detection with all five STAT3 guide RNAs tested (**Fig S5c**). In the LARA model, resting cells are required. Resting memory CD4 T cells were transfected with the combination of two STAT3 guides (guides 3+4) before HIV infection. Non-targeted (NT) guide RNA was used as control. The transfection protocol in resting cells resulted in lower but significant decreased frequencies of STAT3+ cells as measured by flow cytometry (5% full knock-out when compared to the NT guide - p<0.05, n=5 – **Fig S5d**). Additionally, STAT3KO induced on average a decay of 27% at per cell level expression of STAT3 (median fluorescence intensity: MFI – range STAT3KO [MFI 349-486]; range NT [MFI 421-695], p<0.05 – **Fig S5e**). In line with the gene expression results, STAT3 KO show heightened frequencies of dead cells as a consequence of the removal of survival signals downstream of this transcription factor (**Fig S5f,** p <0.0001, Median STAT3hi/dead cells: 34.1%, Median STAT3lo/dead cells: 87.4%) (Left: counter plot gated on STAt3hi x STAt3lo, followed by histogram showing the viability staining in both populations). Knocking out STAT3 prior to infection did not impact the infection rates of these cells with HIV when compared to non-transfected cells or transfected NT (ns: non- significant - **Fig S5g**, n=5). The presence of STAT3hi and STAT3lo cells in resting STAT3-targeted samples allowed us to assess the role of STAT3 in HIV maintenance, and the expression of major proteins of each pathway in the same sample by using a flow-cytometry STAT3 antibody. Gating strategy used for the downstream analysis is shown in **Fig S6 b-d.**

STAT3 knockout led to a 10-fold decay in HIV reservoir, quantified by HIV IntDNA per million CD4 T cells, as compared to NT cells after (FC NT/STAT3KO = 3.0, p<0.01 -**Fig 4c** – evaluation performed 14 days after transfection, 11 days after HIV infection and culture in IL-10 supplemented media). Importantly, cultures of NT transfected cells presented similar levels of HIV IntDNA as non-transfected cultures as both experimental conditions were kept in IL-10 containing media as well (p>0.05, **Fig 4c**). T*hese results confirm the role of IL-10 in inducing latency and HIV persistence*.

### STAT3KO and IL-10 blockade lead to decreased expression of major proteins of key pathways associated with HIV persistence in vivo

Next, we used flow cytometry to validate the expression of representative proteins from each of the pathways associated with pIL-10 and HIV reservoir levels *ex vivo* (survival, memory maintenance, Co-inhibitory receptors (Co-IR), metabolism and Tfh cells) in cells from LARA. These experiments were performed after 11 days of culture in IL-10 containing media in transfected and non-transfected cells. The data analysis was performed in an unbiased way using t-distributed stochastic neighbor embedding (TSNE) or Uniform Manifold Approximation and Projection for Dimension Reduction (UMAP). Briefly, a comparable number of events from each sample was exported and clustered based on the MFI of each marker of interest. Additionally, frequency of cells expressing each marker were quantified. The expression of each marker is provided in both TSNE and UMAP plots (**Fig 4** **d** and **g**, respectively) and in heatmaps (**Fig 4** **e, f** and **h**). In the STAT3KO-targeted cultures, the STAT3hi cells presented significantly increased expression of BCL2 at per cell level (FC hi/lo: 2.48 – p<0.001), increased expression of markers specific to TCM [CCR7 (FC hi/lo: 1.12 – p=0.06) and CD27 (FC hi/lo: 1.36 – p<0.05)] (**Fig 4e**) and higher TCM / TEM ratios (**Fig S7a –** right counter plot shows TCM x TEM in STAT3hi (red) and STAT3 lo (blue) cells); higher per cell level expression of the Co-IR PD1 (PD1- FC hi/lo: 1.19 – p<0.05- **Fig 4e**) and a trend to increased PD1+ cells frequencies (**Fig S7b**) when compared to STAT3lo cells. Importantly, STA3hi cells contributed significantly to the persistence of HIV+ cells (p24- FC hi/lo: 1.26 – p=0.02- **Fig 4e**). *STAT3lo cells (BCL2lo CCR7lo CD27lo PD1lo) presented a significant decrease in the expression of p24, supporting the importance of the survival signal, along with the maintenance of the TCM status and the expression of Co-IRs for HIV persistence*.

We used an independent approach (neutralizing anti-IL-10 mAb) to verify the impact of IL-10 on the modulation of the expression of the key proteins measured above. Non-transfected CD4 T cells were infected and treated with IL-10 [10ng/mL] with or without anti-IL-10 [10ug/ml] for 11 days; expression of BCL2, memory markers, PD1 and HIV protein was measured by flow cytometry. The potential of the anti-IL-10 mAb in blocking IL-10 signaling was shown (**Fig S5b**). A significant decrease in BCL2 expression at per cell level was observed in cell cultures treated with aIL-10, while control cell cultures that received only IL-10 showed increases in BLC2 expression (FC IL-10/aIL-10 1.50, p<0.05 – **Fig 4f**), confirming the transcriptional profiling results (**Fig 3 b-c**). Addition of IL-10 to cell cultures led to an increase in the expression of markers of TCM cells [CCR7 (FC IL-10/aIL-10: 1.99 – p=0.08) and CD27 (IL-10/aIL-10: 2.03 – p=0.004) - **Fig 4f**], as well as an increase in the frequencies of this subset when compared to TEM cells (**Fig S7c**). The per cell expression levels of PD1 were also significantly increased by IL-10 (FC IL-10/aIL-10 1.61 p<0.05 – **Fig 4f**) as well as its frequencies (FC IL-10/aIL-10: 3.54, p<0.01 **– Fig S7d**). Addition of anti-IL-10 decreased the frequencies of PD1+ cells to levels similar to unstimulated cells. Interestingly, our results are also consistent with published results in cancer pre-clinical models(Turnis et al., 2015, Sawant et al., 2019), where IL-10 levels were associated with heightened Co-IR expression. These results validate at protein levels the gene expression profiling (**Fig 3 b-c**) observed ex vivo and specifically induced by IL-10 *in vitro*.

We next monitored the impact of IL-10 signaling on cell metabolism and HIV persistence as suggested by the transcriptional profiling results. IL-10Ra+ cells presented significantly higher levels of glycolysis as monitored by the higher expression of the glucose importer (Glut-1) (FC IL-10Rp/IL-10Rn: 7.00, p<0.0001- **Fig 4g-h, Fig S6d**) additional to higher frequencies of cells expressing Glut-1 (FC IL-10Rp/IL-10Rn 2.79, p<0.05 – **FigS7e**). These Glut-1+ cells were significantly enriched in expression of the HIV-envelope (Env) when compared to cells lacking the engagement to the IL-10 pathway (IL-10Ralo cells) (Env MFI FC IL-10Rp/IL- 10Rn: 1.25, p=0.02 - **Fig 4g-h**). *Collectively, these results validate the ability of IL-10 to regulate the expression of key proteins of the major pathways associated with HIV persistence in vivo. STAT3 KO or neutralization of IL-10 led to decreased expression of the survival protein BCL2, differentiation towards TEM accompanied by downmodulation of PD-1 and decay of immune metabolism resulting in decreased HIV+ cells*.

### IL-10 leads to Tfh differentiation, a major HIV reservoir compartment

Tfh cells are postulated to be the HIV sanctuary in lymph nodes(Fukazawa et al., 2015, Aid et al., 2018). Accordingly, a gene expression signature specific to Tfh cells were found as positive correlate of HIV reservoir in our human cohort and were specifically induced by IL-10 stimulation *in vitro* (**Fig. 3b-c**, respectively). Target genes common to IL-10/STAT3/Bcl6/c-Maf included the major pathways associated with HIV reservoir *in vivo* (survival, Co-IR and Tfh – **Fig 5a**). Of note, 24 hours stimulation of purified CD4 T cells from healthy donors with IL-10, triggered the expression of surface markers and transcription factors known to be features of Tfh cells (CXCR5, PD1, BCL6 and c-MAF)(Crotty, 2014) (FC IL-10/NS: 3.69, p<0.001- **Fig 5b** - Gating strategy **Fig S8a-c).** Addition of anti-IL-10 abrogated the IL-10 induced upregulation of these markers to similar levels of those of unstimulated cells. Interestingly, in another experiment using whole PBMCs and an unbiased UMAP analysis, IL-10 stimulated cells that presented a Tfh profile (STAT3hi, c-Maf hi, BCL6+) were also IL21+ (FC IL-10/aIL-10: 2.07, p<0.01 – **Fig 5c** **-**including inset). IL21 exerts an autocrine impact on Tfh differentiation and helps the formation of the germinal center reaction by triggering B cell proliferation and differentiation(Silver and Hunter, 2008). Importantly, in LARA (non-transfected conditions), cells expressing BCL6 were significantly preferentially infected when compared to their BCL6- counterpart (FC BCL6+/BCL6-: 159.1, p<0.05 - **Fig 5d** – representative counter plot on top show BCL6 vs HIV-p24) mimicking numerous observations showing Tfh as a foyer for HIV persistence(Thacker et al., 2009, Godinho-Santos et al., 2020, Cai et al., 2019). In chronically treated HIV-infected individuals, Tfh cells become a reservoir sanctuary in the BCF (F), being enriched in HIV IntDNA (**Fig 5e**). *These results show that IL-10 by inducing Tfh differentiation in BCF could provide an important source of targets for HIV infection and viral persistence*.

**Fig 5.**
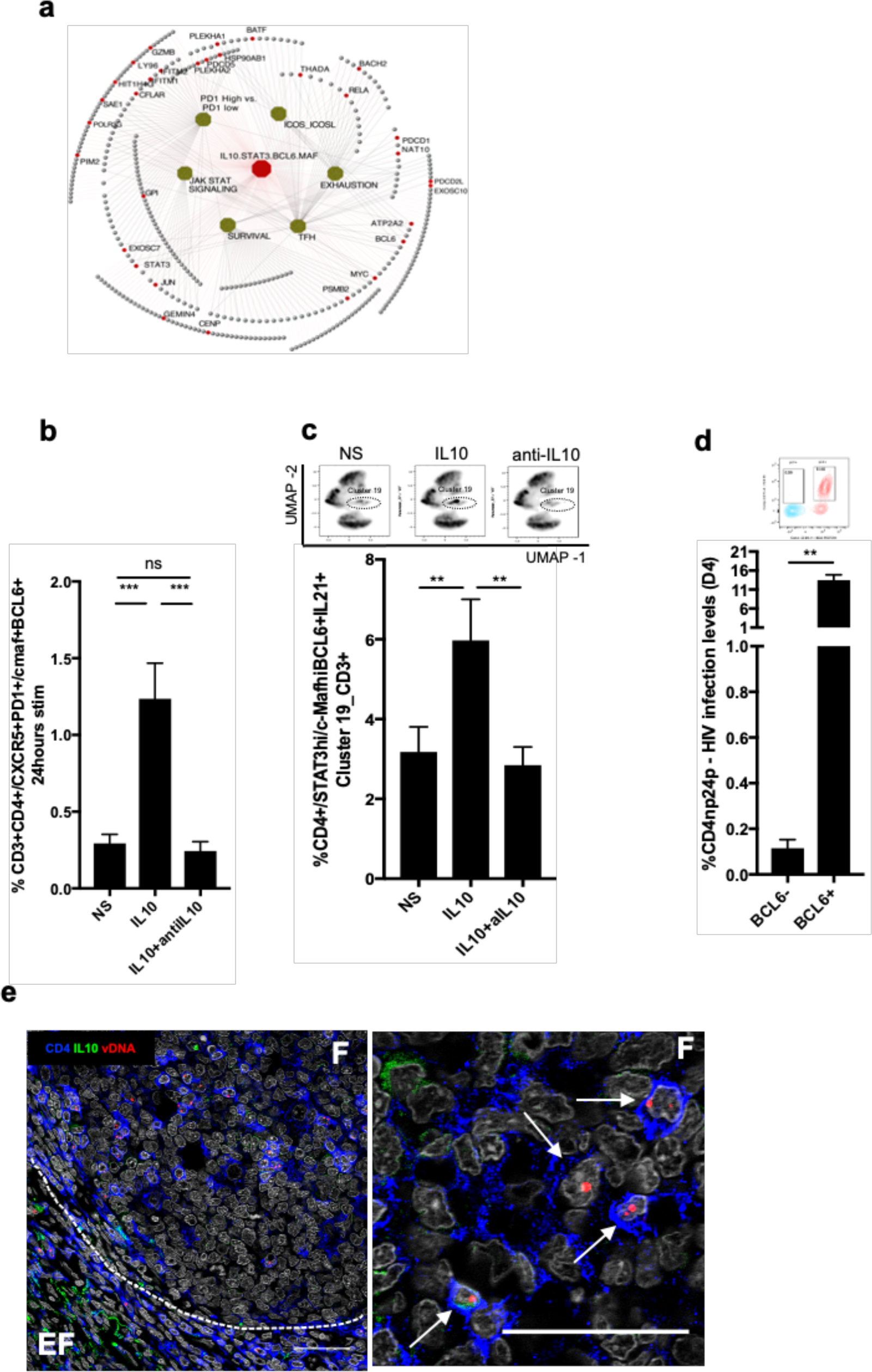
IL-10 leads to Tfh differentiation, a major HIV reservoir compartment. **a)** IL10/STAT3/BCL6/c-maf (red node) LEGs (open dots) are shared and are part of the major pathways associated to pIL-10 levels and HIV IntDNA ex vivo (green nodes). Red-full dots, represent important genes for the function of each pathway. Cytoscape was used to generate the gene network. **b)** PBMCs from healthy donors were treated with IL-10 (10 ng/mL) +/- anti-IL10 mAb (10ug/mL) or left unstimulated in complete media for 24 hours. Brefeldin was added for extra 6 hours in culture (total 30hours). The expression of Tfh markers (CXCR5, PD1), its major transcription factors (c-Maf and Bcl6) as well as the production of IL-21 were evaluated by flow cytometry. **c)** After UMAP analysis, Cluster 19 was specifically induced by IL-10. On top, density plots show the modulation of cluster 19 in the different conditions. This cluster is STAT3hi/c-Mafhi, BCL6+ and IL21+ (counter plots shown in **Fig S8c)**. **d)** In the LARA model, 4 days after infection, BCL6+ cells are significantly preferentially infected when compared to its BCL6- counterpart; top: representative dot plot, bottom: quantification of CD4-p24+ cells in BCL6- or BCL6+ T cells; **e)** Immuno-histochemetry in LNs from HIV aviremics individuals. Follicular (F) and Extrafollicular (EF) are shown (left image: white dotted line). In the follicles, CD4+ T cells (blue) are enriched in HIV reservoir (HIV DNA – red and white arrows). IL10+ cells are shown (green).

## DISCUSSION

Persistent latent infection requires that HIV infected cells endure over time. Herein, we identified (*in vivo*) and validated (*in vitro*) our hypothesis that IL-10 plays critical role for the maintenance of the HIV reservoir. We have demonstrated that IL-10 enhances i) cell survival, ii) upregulates the expression of the co-inhibitory receptor PD-1 (several Co-IRs are part of the gene signature, such as Lag3, TIGIT, CTLA4), which have been previously shown to be involved in the establishment of HIV latency and immune dysfunction(Trautmann et al., 2006, Tzeng et al., 2012, Peng et al., 2008); iii) prevents the differentiation of quiescent TCM to the TEM subset, which we have shown earlier to favor HIV replication(Wonderlich et al., 2019); iv) triggers a *bona fide* metabolic status associated with long lasting cells; and finally, v) we have shown that IL-10 is a potent regulator of Tfh differentiation. *The role of IL-10 in HIV persistence is supported by our findings where knocking out STAT3, the transcription factor downstream of IL-10/IL-10R engagement(Donnelly et al., 1999), led to decreased frequencies of cells with HIV DNA as a consequence of the downregulation of the key pathways sustaining reservoir survival, latency and longevity (metabolically active TCM cells)*. We have shown that IL-10 is produced by several cellular subsets including innate immune cells, B cells and T cells i.e most probably Tr1(Zhang and Kuchroo, 2019) or Tfr cells(Laidlaw et al., 2017) in all 3 anatomical sites in the lymph nodes. IL-10 expression is controlled by several transcription factors(Kubo and Motomura, 2012, Zhang and Kuchroo, 2019) that can be triggered by TLRs as shown by our results (NFKB1, EOMES, STAT1) and/or by TCR/CD28 engagement (NFKB1, EOMES, STAT1)(Zhang and Kuchroo, 2019). Microbial products can also trigger B cells and innate immune cells to produce IL-10(Boonstra et al., 2006, Saemann et al., 2000, Liu et al., 2014), which in turn, will promote the differentiation of IL-10 producing Tr1 cells(Asseman and Powrie, 1998). Given that we and others have demonstrated that gastrointestinal tract damage and dysfunction, and microbial translocation persists in individuals on long-term ART(Somsouk et al., 2015), there is a continual systemic source of stimulation that could result in high levels of IL-10 driving viral persistence. Further *in vivo* models suggest a role for the gut flora in triggering IL- 10 and regulating homeostatic interactions with commensal microorganisms as IL- 10 dependent immunopathology is reversed under germ-free conditions(Sellon et al., 1998). While our results do not discriminate between cells that bind IL-10 and cells that produce IL-10 we provide herein statistically significant evidence that the proximity of IL-10+ cells to HIV DNA+ cells in the LN tissues further supports the notion that IL-10 is important for HIV persistence *in vivo* in tissues. The co- localization of pSTAT3+ and vDNA+ cells to IL-10+ cells, and the increased frequencies of vDNA+ cells in the vicinity of IL10+ cells further implicates this cytokine in promoting HIV persistence in lymph nodes. IL-10 induces latency by downregulating the expression of several transcription factors that are involved in promoting HIV transcription including NFK-b, C/EBPb, NFAT, Ets/PU.1(Schiralli Lester and Henderson, 2012), or by promoting the expression of TFs that can repress HIV LTR activity (PRDM-1)(Kaczmarek Michaels et al., 2015). In the periphery, IL-10 also plays a major role in HIV persistence as demonstrated by the fact that IL-10 was unique among all cytokines that signal through STAT3, including IL-6 and IL-21, to be associated with HIV reservoir size. Differences in signaling and transcriptional activity between IL-10 and IL-6 have been previously reported. Expression of SOCS3 upon IL-10R engagement does not lead to IL-10 receptor degradation, keeping IL-10 signaling constant(Niemand et al., 2003).

IL-10 signaling enhances the survival of different immune and non-immune cell subsets(Zhou et al., 2001, Levy and Brouet, 1994, Todaro et al., 2006, Boyd et al., 2003) including B cells, which are critical for the maintenance of Tfh numbers(Cubas et al., 2013). While many studies associate IL-10 induced cell survival to BCL2 expression, here we show that the increased survival involves the upregulation of several anti-apoptotic molecules (Bcl2, BclX-l, Pim1, BIRC5) downstream of IL-10 signaling. Enhanced survival could have a direct impact on HIV persistence by enhancing the dissemination of infectious HIV virions to bystander cells in sites where antiviral drugs show poor levels of penetrance(Gavegnano et al., 2017). Additional to endorsing cell survival, homeostatic proliferation(Virgilio and Collins, 2020) and/or clonal expansion(Liu et al., 2020) could further contribute to increased size of the HIV reservoir. Importantly, the usage of an FDA-approved BCL2 antagonist, venetoclax, during HIV reactivation, led to cell death and HIV reservoir decay(Cummins et al., 2017), we also observed that IL-10 signaling blockade led to HIV decay.

Our data shows that the heightened levels of pIL-10 is associated with increased expression of Co-IRs, including PD-1, which we and others have shown to trigger T cell quiescence and HIV latency(Evans et al., 2018, Porichis and Kaufmann, 2012, Fromentin et al., 2019). Early upon infection, cellular activation leads to PD- 1 expression as part of the cascade and HIV preferentially replicates in PD-1+ CD4 T cells(Vollbrecht et al., 2010). However, the binding of PD-1 and other Co-IRs to its ligands triggers T cell quiescence(Fromentin et al., 2019); additionally, the expression of PD-1 blocks the differentiation of TCM cells to TEM(Trautmann et al., 2006, Breton et al., 2013, Kulpa et al., 2013). This mechanism provides support for the association between IL-10, PD-1 and frequencies of cells with HIV integrated DNA. Interestingly, the expression of Co-IRs was shown to lead to increased IL-10 production(Dong et al., 1999) upon ligand binding. Of note, we have previously shown that PD-1 stimulation can induce production of IL-10 by myeloid cells and suppression of antiviral immunity(Said et al., 2010). IL-10 triggers pathways associated with cell quiescence, and further impedes effector cell differentiation by upregulating transcriptional factors (TFs) associated with maintenance of TCM (TCF7, Notch, FOXO1 and Foxp1(Tang et al., 2012)), and by downregulating TFs critical for effector function (NFAT, NFKb). Furthermore, as mentioned previously, the upregulation of PD-1 by IL-10 is associated with blockade of T cell memory differentiation and their accumulation as TCM(Trautmann et al., 2006, Breton et al., 2013, Kulpa et al., 2013); this would contribute to the increased accumulation of these cells that are the long-lasting HIV reservoir. Heightened pIL-10 levels have been associated with poor effector function and absence of viral clearance in chronic viral infections(Tian et al., 2016) most probably due to the lack of differentiation of TCM to cells endowed with antiviral effector activity. Relevant to this hypothesis, T cells from IL-10-/- mice exhibited a highly activated phenotype, expressed antiviral cytokines and promoted degranulation in response to cognate Ag-encounter *ex vivo* and were oligoclonal suggesting an ongoing immune response(Jones et al., 2010). In addition to preventing the differentiation of TCM to TEM, IL-10 promotes survival of all memory cell subsets including TEM, which was shown to have the highest levels of inducible HIV(Kulpa et al., 2019). In our experimental model, IL-10 blockade *in vitro,* removed the survival signal, additional to promoting T cell activation, by downmodulating Co-IRs, and differentiation towards TEM. Importantly, IL-10 helps to fuel cell longevity by promoting a balanced immune metabolism. Much is known about the impact of IL-10 on the metabolism of innate immune cells as it promotes oxidative phosphorylation in macrophages(Ip et al., 2017). IL-10 induces mitophagy that eliminates dysfunctional mitochondria characterized by low membrane potential and a high level of reactive oxygen species(Ip et al., 2017). In the absence of IL-10 signaling, macrophages accumulate damaged mitochondria(Ip et al., 2017). *Herein, we show that IL-10 triggers enhanced cell metabolism by promoting glycolysis*.

The impact of IL-10 on Tfh differentiation is significant, and to the best of our knowledge this is the first time that these findings are reported. Previous results have shown that IL-10 levels are associated with poor development of functionally mature memory Th1 cells while being positively correlated to increased frequencies of Tfh cells during acute LCMV infection(Tian et al., 2016). Tfh cells are known to harbor HIV intact provirus and can be a source of replication competent virus(Perreau et al., 2013) under ART interruption. Tfh cells provide help for the differentiation and maturation survival of GC B cells(Crotty, 2019), that are known to produce IL-10(Burdin et al., 1997). Additionally, in the GC environment, the IL-10 production is boosted by IL-21, produced by Tfh cells, and the phosphorylation of STAT3 further enhances IL-10 production(Banko et al., 2017). A complex network of TFs, including BCL-6, interferon regulatory factor (IRF4), c-Maf, and BATF, promotes Tfh differentiation while inhibiting alternate CD4 T cell differentiation pathways(Aid et al., 2018), all upregulated at gene and/or protein level in our study and associated with the size of the HIV reservoir. Indeed, Tfh cells were demonstrated to be both productively (i.e., vRNA^+^) and latently (i.e., vDNA) infected at higher frequencies(Perreau et al., 2013, Lindqvist et al., 2012) than non-Tfh cells. Tfh cell accumulate in chronic infection(Maceiras et al., 2017) and this could be related to the higher levels of IL-10 in LNs of HIV-aviremic individuals as shown by our results. Tfh cells can be characterized by the expression of PD1, that is also upregulated by IL-10. Additionally, we have observed BCL2L12 known to promote T cell survival is also included in the Tfh signature. In the current study, we have shown that IL-10+ and vDNA+ cells are colocalized in the GC area, indicating that aside of promoting the differentiation and survival of the Tfh cells, IL-10 is consequently contributing to the survival of the HIV reservoir in this important compartment.

Pre-clinical interventions using anti-IL-10 have been performed in several studies and despite some concerns regarding safety, have led to positive outcomes. The blockade of IL-10 in the LCMV model led to the inhibition of viral persistence and enhanced T-cell functions(Ejrnaes et al., 2006, Brockman et al., 2009). IL-10 blockade, in addition to clear chronic viral infections in animal models, also improved cytotoxic T cell responses in immunization models(Pitt et al., 2012). In HIV infection, IL-10Ra blockade resulted in markedly increased secretion of IFN-g by CD4+ T cells(Saeidi et al., 2018) as well restored the polyfunctionality of HCV specific T cells(Wilson and Brooks, 2011, Rigopoulou et al., 2005). Dual IL- 10R/PD1 blockade further enhanced T cell activity, suggesting that IL-10 and PD- 1/PD-L1 are functional through distinct pathways to suppress T cell activity during persistent viral infection(Brooks et al., 2008). Of note, pegylated IL-10 has been shown to enhance CD8 T cell function in cancer patients(Naing et al., 2018, Mumm and Oft, 2013) in sharp contrast to several other studies, where IL-10 negatively impact T cell function(Kahan and Zajac, 2019). Our results show that an anti-IL-10 antibody(L., 2005), was effective in down-regulating *in vitro* the expression of proteins which are part of major pathways associated *in vivo* to HIV persistence, leading to HIV decay in vitro. *Altogether our results feature IL-10 blockade as a promising and powerful strategy to revert immune-suppression and to impede on HIV reservoir maintenance leading to a HIV cure*.

## STAR METHODS

### EX VIVO

#### Cohort

HIV-Aviremic individuals (n= 42) and HIV negative healthy controls (n=10) (**Table S1**) signed informed consent approved by the Royal Victoria Hospital and the CRCHUM Hospital Institutional Review Boards. Plasma, Peripheral blood mononuclear cells (PBMCs) and whole blood paxgene tubes were collected. Plasma viral load was evaluated with the Amplicor HIV-1 Monitor Ultrasensitive Method (Roche)(Sun et al., 1998).

#### Isolation of CD4+ T cells

PBMCs were isolated from blood samples or leukapheresis products by Ficoll Hypaque density gradient centrifugation(Fuss et al., 2009). Total CD4+ T cells were isolated from PBMCs by negative magnetic selection (Stemcell technologies). The purity of enriched CD4+ T cells was generally greater than 95%, as assessed by flow cytometry (data not shown).

#### Plasma cytokines quantification

##### Mutiplex ELISA (Mesoscale)

U-PLEX assay (Meso Scale MULTI-ARRAY Technology) commercially available by Meso Scale Discovery (MSD) was used for plasma cytokine detection. This technology allows the evaluation of multiplexed biomarkers by using custom made U-PLEX sandwich antibodies with a SULFO-TAG™ conjugated antibody and next generation of electrochemiluminescence (ECL) detection. The assay was performed according to the manufacturer’s instructions (https://www.mesoscale.com/en/technical_resources/technical_literature/techncal_notes_search). In summary, 25μL of plasma from each donor was combined with the biotinylated antibody plus the assigned linker and the SULFO-TAG™ conjugated detection antibody; in parallel a multi-analyte calibrator standard was prepared by doing 4-fold serial dilutions. Both samples and calibrators were mixed with the Read buffer and loaded in a 10-spot U-PLEX plate, which was read by the MESO QuickPlex SQ 120. The plasma cytokines values (pg/mL) were extrapolated from the standard curve of each specific analyte. Plasma cytokines levels were analyzed to test for differences between HIV-aviremics and Health Controls (HC). Unpaired non-parametric Mann-Whitney T-test was performed. Negative log 10 (-log_10_) values of the cytokines was plotted against log2 of the fold change (FC) between HIV aviremics and HIV negative healthy controls. Linear regression and Spearman correlation were performed against HIV IntDNA measures. Negative log 10 (-log_10_) values of HIV IntDNA was plotted against Spearman rho values HIV aviremics.

### Integrated HIV DNA quantification

CD4+ T cells isolated from aviremic donors were digested and cell lysates were directly used in a nested Alu PCR to quantify both integrated HIV DNA and CD3 gene copy numbers, as previously described(Vandergeeten et al., 2014).

### Microarray

Gene signature profile was assessed using Illumina Human version 4 beadchips (Illumina) at 58 °C for 20 h. RNA was isolated using the Rneasy micro kit (Qiagen) and the quantity and quality of the RNA were confirmed using a NanoDrop 2000c (Thermo Fisher Scientific) and an Experion Electrophoresis System. Samples (50 ng) were amplified using Illumina TotalPrep RNA amplification kits (Ambion) as per manufacture instructions. The chips were scanned using Illumina’s iSCAN and quantified using Genome Studio (Illumina). Raw beadchips intensities were quantile-normalized and log2-transformed.

### Multiparameter confocal imaging

Confocal imaging was performed with formalin fixed paraffin embedded (FFPE) lymph node sections prepared at a ∼5 μm thickness. Tissue sections were deparaffinized by bathing in xylene and serial ethanol dilutions. Antigen retrieval was performed at 110°C for 15 minutes using Borg Decloaker RTU (Biocare Medical). Tissue sections were then blocked, permeabilized for 1 hour at room temperature and stained with the following primary and conjugated antibodies: anti-CD20 eFluor 615 (clone L26), anti-Ki67 Brilliant Violet 421 (clone B56), anti-CD4 Alexa Fluor 488 (Goat polyclonal IgG, FAB8165G, R&D systems), anti-IL-10 (rabbit polyclonal, ab34843, Abcam) and either anti FoxP3-Alexa Fluor 647 (clone 206D) or anti CD68 (clone KP-1) and CD163- Alexa Fluor 647 (clone EDHu-1) depending on the panel. The nuclear stain Jojo-1 (Life Technologies) was also used to delineate individual cells. Stainings were carried out consecutively with the primary antibodies being added first and incubated overnight at 4° C, followed by staining with the appropriate secondary antibody which –depending on the panel - was either Alexa Fluor 546 goat anti-rabbit IgG for IL-10 (ThermoFisher Scientific, A11010) or Alexa Fluor 488 goat anti-mouse IgG1 for CD68 (ThermoFisher Scientific, A21121). Conjugated antibody stainings were performed for 2 hours at room temperature, after which sections were stained with JoJo-1 for nucleus identification. Images were acquired at a 512 x 512 pixel density using a 40x objective (NA 1.3) on a Nikon confocal system running NIS-elements AR. Fluorophore spectral spillover was corrected through live spectral un-mixing (NIKON).

### Histo-cytometry and imaging analysis

Post-acquisition analysis was performed using the Imaris software (Bitplane, version 8.4). Histo-cytometry was performed as previously described(Gerner et al., 2012, Petrovas et al., 2017). Briefly, dimensional imaging datasets were segmented based on their nuclear staining signal and average voxel intensities for all channels were extrapolated in Imaris after iso-surface generation. Data were then exported to Microsoft Excel, concatenated into a single comma separated values (csv) format and imported into FlowJo version 10 for further analysis. The results of the histocytometry quantification were expressed as average frequencies.

### Next-generation HIV DNA in situ hybridization and immunofluorescent detection for confocal phenotypic analysis

The next-generation *in situ* hybridization method DNAscope(Deleage et al., 2016) and multiplex immunofluorescence staining were performed on 5 μm thick sections from FFPE lymph node biopsies from HIV-1 infected, cART suppressed participants from Emory University and Case-Western Reserve University cohorts, along with FFPE ACH-2 HIV-1 positive controls [obtained through the NIH AIDS Reagent Program, Division of AIDS, NIAID, NIH: ACH-2 Cells from Dr. Thomas Folks (cat# 349)](Clouse et al., 1989, Folks et al., 1989) and human HIV-uninfected lymph node negative controls from the OHSU Biolibrary. DNAscope was performed according to Deleage *et al*(Deleage et al., 2016) with some modifications. Briefly, slides were deparaffinized by heating at 60°C for 1 hour, followed by two 5-minute xylene incubation and two 1-minute 100% ethanol incubations. Slides were then kept in double distilled water for 1 hour before heat induced epitope retrieval using the ACD Target Retrieval Buffer and the Biocare NxGen Decloaking Chamber that was set to 110°C for 15 min. After the Decloaking Chamber was cooled to 95°C, the slides were removed from the chamber and cooled at room temperature for a further 15 minutes. Slides were then rinsed twice in double distilled water and tissues were kept wet throughout the rest of the protocol. Protease treatment was omitted to improve antibody epitope preservation and staining. Slides were incubated for 10 minutes at room temperature with 3% Hydrogen peroxide diluted in PBS to inactivate endogenous peroxidases. The HIV- 1 Clade B sense probe (ACDbio cat. No. 425531) was added to the tissue for a 14-hour incubation at 40°C in a HybEz oven. The ACDBio RNAscope 2.5 Brown kit was used to amplify vDNA signal according to manufacturer’s recommendations, with the exception that wash steps were performed with 0.5X RNAscope wash buffer. In addition, after Amp 5 and 6, slides were washed with 1X TBS-Tween-20 (0.05% v/v) instead of ACD wash buffer. Viral DNA fluorescence signal was developed using the ThermoFisher Alexa Fluor (AF)647 Tyramide Reagent followed by boiling in ACDBio Target Retrieval buffer for 10 minutes at 95-100°C to inactivate the HRP. Immunofluorescence antibody staining was performed using a rabbit anti-Human IL-10 antibody (Abcam ab-34843), a rabbit anti-pSTAT3 antibody (Cell Signaling 9145) and a mouse anti-CD20 antibody (Dako M0755). The anti-IL-10 antibody was labelled using an HRP-conjugated polymer anti-rabbit secondary antibody (GBI Labs D13-110) and developed with AF568 Tyramide reagent (ThermoFisher), followed stripping of the primary rabbit antibody and inactivation of the HRP by boiling in ACDBio Target Retrieval buffer for 10 minutes at 95-100°C. Subsequent anti-pSTAT3 and anti- CD20 staining were labelled with AF 750 conjugated anti-rabbit and AF488 conjugated anti-mouse secondary antibodies (ThermoFisher) respectively. Tissues were counterstained with DAPI and cover slipped with #1.5 GOLD SEAL cover glass (EMS) using Prolong® Gold reagent (ThermoFisher).

### Quantitative image analysis

To quantify the number of vDNA+, IL-10+ and/or pSTAT3+ cells on stained tissues, whole-slide high-resolution fluorescent scans were performed at 20X using the Zeiss AxioScan Z.1 slide scanner. DAPI, AF488, AF568, Cy5 (For AF647) and Cy7 (For AF750) channels were used to acquire images. The exposure time for image acquisition was between 4 and 150 ms. Multi-spectral images were analyzed using the HALO 2.3 platform (Indica Labs) using the following analysis methods: For HIV-1 vDNA, cells with clear punctate dots were quantified using the module FISH v1.1. Thresholds for spot size and intensity were standardized against the vDNA signal in ACH-2 cells, which has at least one integrated provirus per cell. Total cell count per tissue was also performed using this module. For the quantification of the total number of pSTAT3+ and IL-10+ cells per tissue, the FISH v1.1 module was also used. The Nearest Neighbor and Proximity Analysis were performed with the Spatial Analysis plot function of HALO, using the object data derived from the previous individual analyses discussed above. Manual curation was performed to confirm the accurate quantification of vDNA+ cells and their individual IL-10 and pSTAT3 status.

### IN VITRO

#### Primary CD4+ T Cell Isolation and Culture

Briefly, primary human CD4+ T cells from healthy donors were isolated from donated leukoreduction chambers. Peripheral blood mononuclear cells (PBMCs) were isolated by Ficoll centrifugation using SepMate tubes (STEMCELL, per manufacturer’s instructions). Bulk CD4+ T cells were subsequently isolated from PBMCs by magnetic negative selection using an EasySep Human CD4+ T Cell Isolation Kit (STEMCELL, per manufacturer’s instructions). Alternately, memory CD4+ T cells were isolated from PBMCs by magnetic negative selection using an EasySep™ Human Memory CD4+ T Cell Enrichment Kit (STEMCELL, per manufacturer’s instructions). Isolated CD4+ T cells were suspended in complete Roswell Park Memorial Institute (RPMI), consisting of RPMI-1640 (Sigma) supplemented with 5mM 4-(2- hydroxyethyl)-1-piperazineethanesulfonic acid (HEPES, Corning), 2mM Glutamine (UCSF Cell Culture Facility), 50μg/mL penicillin/streptomycin (P/S, Corning), 5mM sodium pyruvate (Corning), and 10% fetal bovine serum (FBS, Gibco).

#### IL-10 stimulation in vitro

Viable frozen primary human PBMCs cells from healthy donors were thawed, counted and rested for 2 hours at 37C, 5% CO2 at a concentration of 2 million PBMCs/mL in vented cap bottles in complete Roswell Park Memorial Institute (RPMI), consisting of RPMI-1640 (Sigma) supplemented with 5mM 4-(2- hydroxyethyl)-1-piperazineethanesulfonic acid (HEPES, Corning), 2mM Glutamine (UCSF Cell Culture Facility), 50μg/mL penicillin/streptomycin (P/S, Corning), 5mM sodium pyruvate (Corning), and 10% fetal bovine serum (FBS, Gibco). Upon resting, 1 million PBMCs were transferred to 48-well plate left unstimulated or were stimulated with IL-10 [10ng/mL] plus or minus anti-IL-10 [10ug/mL] for 24 hours. Brefeldin (1:1000) was added for extra 6 hours in culture (total 30hours). The expression of Tfh markers (CXCR5, PD1, c-Maf and Bcl6 and IL21) were evaluated by flow cytometry.

To evaluate IL-10 signaling through phosphorylation of STAT3 (pSTAT3), different doses of anti-IL-10 was used [0.1-100ug/mL] in addition to IL-10 [10ng/mL]. pSTAT3 MFI was evaluated after 30 minutes of stimulation by flow cytometry.

#### Microarray for IL-10 induced specific signatures in CD4 T cells

CD4 T cells were isolated from PBMCs from healthy donors (n= 6) were cultured for 12 hours with 10ng/mL of rIL-10 or left unstimulated. Microarray was performed as for the *ex vivo* experiments.

### STAT3KO

#### RNP Production

Detailed protocols for RNP production and primary CD4+ T cell editing have been previously published(Hultquist et al., 2019). Briefly, lyophilized crRNA and tracrRNA (Dharmacon) were suspended at a concentration of 160 µM in 10 mM Tris-HCL (7.4 pH) with 150 mM KCl. 5µL of 160µM crRNA was mixed with 5µL of 160µM tracrRNA and incubated for 30 min at 37°C. The gRNA:tracrRNA complexes were then mixed gently with 10µL of 40µM Cas9 (UC-Berkeley Macrolab) to form Cas9 ribonucleoproteins (RNPs). Five 3.5µL aliquots were frozen in Lo-Bind 96-well V-bottom plates (E&K Scientific) at -80°C until use.

#### Editing of Resting Memory CD4+ T Cells

Each reaction consisted of 30x10^6 CD4 T cells, 3.5 µL RNP, 1 µL Alt-R Cas9 Electroporation Enhancer (100 µM, IDT) and 20 µL electroporation buffer. Immediately after isolation, resting memory CD4+ T cells were suspended and counted. RNPs were thawed and allowed to come to room-temperature. One microliter of Alt-R Cas9 Electroporation Enhancer was added to each RNP mixture with gentle mixing. Immediately prior to electroporation, cells were centrifuged at 400xg for 5 minutes, supernatant was removed by aspiration, and the pellet was resuspended in 20 µL of room-temperature P2 electroporation buffer (Lonza) per reaction. Twenty microliters of cell suspension were then gently mixed with each RNP mixture and aliquoted into an electroporation cuvette for nucleofection with the 4D 96-well shuttle unit (Lonza) using pulse code EH-100. Immediately after electroporation, 80 µL of pre-warmed media without IL-2 was added to each well and cells were allowed to rest for at least one hour in a 37°C cell culture incubator. Subsequently cells were moved to vented-bottomed culture bottles pre-filled with warm complete media with IL-2 at 40 IU/mL (for a final concentration of 20 IU/mL). Cells were cultured at 37°C / 5% CO2 in a dark, humidified cell culture incubator for 4 days to allow for gene knock- out and protein clearance, with additional media added on day 2. To check STAT3 knock-out efficiency, 50 µL of mixed culture was removed to a centrifuge tube. Cells were staining for total STAT3 monoclonal antibody by flow cytometry. For the LARA experiments a combination of guides 3 and 4 from Dharmacon was used (CM-003544-03-0020, CM-003544-04-0020).

#### LARA (Latency and Reversion assay)

Briefly, on day 0 memory CD4 T cells were isolated. T cells were immediately knocked out for STAT3 or NT and kept in IL-10 [10ng/mL] containing media or non-transfected and cultivated in cRPMI for 2-3 days. On Day 4, cells were HIV infected by spinoculcation with the full-length replication competent HIV clone 89.6. Immediately after spinoculation, cells were resuspended in IL-2 [30U/mL] and Saquinavir [5uM]. Infected cells were incubate for an additional 2-3 days before being introduced into latency culture conditions. For the transfected conditions latency media was composed of antiretroviral cocktail (ARV) of 100 nM efavirenz, 200 nM raltegravir, and 5 μM saquinavir (NIH AIDS Reagent Program; 4624, 11680, and 4658), IL7 [40ng/mL] and IL-10 [10ng/mL]. Non-transfected conditions were cultured in ARV+ IL-7, ARV+ IL-7+IL- 10, ARV+ IL-7+IL-10+aIL-10 [10ug/mL] in order to evaluate the role of IL-10 on latency induction. Gold standard latency inducer media was ARV+ IL-7+TGFb1 [20ng/mL] in media composed 50% of H80 supernatant culture(Kulpa et al., 2019). Cells were maintained in this media for 11 days. Half media change was performed every 3-4 days. HIV protein expression decay as the evaluation of other major proteins, of the IL-10 modulated pathways were accessed by flow cytometry. Ten thousand cells were saved at the last day of latency for HIV IntDNA evaluation in the IL-10 containing media conditions.

#### Cell Preparation and flow cytometry

PBMCs were prepared from whole blood by ficoll-hypaque density sedimentation and cryopreserved in 10% dimethyl sulfoxide and 90% FBS until thawing for phenotypic analysis. Panels to evaluate Survival, Co-IRs, memory subsets, Immune metabolism, Tfh markers and HIV protein expression were performed. Markers were combined in different panels to address the above questions. Antibodies summarized in **Table S7** were used. Flow cytometry data were analyzed to test for differences among the different stim conditions. All antibodies were properly titrated. The cells were all surface stained for 20 minutes in the dark at room temperature, washed, fixed and permeabilized using the eBioscience™ Foxp3 / Transcription Factor Staining Buffer Set (Cat# 00- 5523-00), as per manufacture instructions. Intracellular staining was performed in Perm-Wash provided by the kit for 45 minutes at 4°C. Samples were then washed and re-suspended in staining buffer for acquisition. Samples were acquired on LSRII flow cytometer (Becton Dickinson, San Jose, CA) and ∼500,000 live-gated events were collected. Data were analyzed using Flow-Jo software (TreeStar, Ashland, OR). A lymphocyte gate based on FSC-A and SSC-A was defined. Single cells were then selected using FSC-A x FCS-H gate. Live cells were gated and successive gates to define T cell populations of interest. TSNE analysis (t-distributed stochastic neighbor embedding) or UMAP analysis (Uniform Manifold Approximation and Projection for Dimension Reduction) were performed in live single cells for unbiased evaluation of the distribution of the key markers of the major pathways identifies by gene expression *ex vivo* and *in vitro*.

#### Reagents, antibodies and viruses

IL-10 from Peprotech was used at 10ng/mL; IL-7 from R&D Systems was used at 40ng/mL; TGFb1 from Peprotech was used at 20 ng/mL; anti-IL-10 was produced and characterized by D. Gorman(L., 2005) specifically for this project, dose-response curve was performed evaluating blockage of STAT3 phosphorylation (**Fig S5b**) and 10ug/mL was used for the downstream analysis. Flow antibodies were acquired from BD, Biolegend, MedMab, Beckman Coulter, eBioscience and titrated for best performance in each flow panel (**Table S7**). Viruses were acquired from NIH reagents program (p89.6) and were produced by the virus core facility at CWRU.

### IN SILICO

#### Microarray Analysis

Gene array was performed in PBMCs from HIV aviremic individuals. For each gene, a linear regression model with pIL-10 or HIV intDNA measurements in aviremic individuals as an independent variable and gene expression as a dependent variable was fit using the R package LIMMA. Genes that correlated with both outcomes using a Benjamini– Hochberg corrected p value of 0.05 were selected. Gene sets enrichment analysis (GSEA) in ex vivo samples revealed significant and positive correlation of gene signature with IntDNA and pIL- 10 levels. Gene signatures were compiled from MSigDB C2 and C7 and other gene sets available in the literature. The LIMMA package was used to fit a linear regression model to each probe (log2 expression) to respectively IntDNA and pIL- 10 levels. Genes that correlated to the outcomes were sorted from high to low using their coefficient correlation for each regression and then submitted to GSEA using the pre-Ranked list option. Signatures significantly enriched in these lists with an FDR cut-off of 5% were selected. GSEA normalized enrichment scores were plotted using pheatmap R package. Color gradient from blue to red depicts the normalized enrichment score ranging from decreased (blue) to increased (red). For the *in vitro* IL-10 induced gene signature the LIMMA R package was used to fit linear regression model with the log2 gene expression as dependent variable and the IL-10 stimulated and IL-10 unstimulated CD4 T cells as independent variables in order to identify genes differentially expressed between CD4 T cells stimulated by IL-10 at 12 hours compared to unstimulated CD4 T cells or genes correlated to plasma IL-10 levels and HIV integrated DNA from ART suppressed (aviremic) samples. A moderated *t* test was used to assess the statistical significance of the association between gene expression and the groups of interest. The p value was adjusted for multiple testing using Benjamini and Hochberg correction method.

To identify pathways enriched in IL-10 stimulated genes at 12 hours and pathways enriched in genes correlated with plasma IL-10 levels and HIV integrated DNA in ART suppressed individuals, we use The GSEA Java desktop program was downloaded from “http://www.broadinstitute.org/gsea/index.jsp[http://www.broadinstitute.org/gsea/index.jsp[http://www.broadinstitute.org/gsea/index.jsp]” and the default parameters of GSEA preranked option with the following parameters: gene list ranked by their fold; number of permutations: 1000; enrichment statistic: weighted; seed for permutation: time, ignore genesets with less than 10 genes or more than 5000 genes; and the following pathways databases: Molecular Signatures Database (version 5.1) hallmark genesets^46^, canonical pathways (module C2.CP), transcription factor targets (module C3.TFT). Specific transcription factors target signatures including STAT3, BCL6 and MAF were extracted from pscan database (http://www.beaconlab.it/ pscan) and the we used a T follicular helper signature as defined in Aid *et al*(Aid et al., 2018).

For pathways enrichment analyses, we applied a 5% cutoff on the false discovery rate (FDR) and a corrected p value of 0.05 on all the differential expression and the regression analyses. Genes network analyses and representation were performed using GeneMania (http://genemania.org) and DyNet Analyzer application under Cytoscape 3.6.0 ( https://cytoscape.org).

#### Pathways enrichment analysis

Gene set enrichment analysis was performed using GSEA and a compiled set of pathways from public data bases including MSigDB version 5.1 (http://software.broadinstitute.org/gsea/msigdb/) and blood cell marker signatures. To test for the enrichment of IL-10 and PD-1 signaling we used in-house signatures (unpublished data). The GSEA Java desktop program was downloaded from the Broad Institute (http://www.broadinstitute.org/gsea/index.jsp) and used with GSEA Pre-Ranked module parameters (number of permutations: 1,000; enrichment statistic: weighted; seed for permutation: 111; 10 ≤ gene set size ≤ 5,000). We used Dynet Analyzer application implemented in Cytoscape version 3.6.0 to generate gene interacting networks to highlight overlapping genes between the different enriched modules.

#### Statistics

Data sets were tested for a Gaussian distribution using the D’Agostino-Pearson omnibus normality test. Dependent upon the determination of data normality, correlations against plasma IL-10 concentrations were conducted using either a Pearson or nonparametric Spearman analyses. Longitudinal analyses comparing the concentration of plasma IL-10, the %IL-10^+^ cells in LN by anatomical site, and vDNA^+^ cells in LN were calculated using either a Mann-Whitney U test or a Wilcoxon matched-pairs signed rank test dependent upon the replicate pairing and Gaussian distribution of the data. Analyses of the *in vitro* stimulations were conducted using a one-way ANOVA with matching and multiple comparisons with a Tukey correction. All statistical tests were two-sided and not adjusted for multiple comparisons. The shorthand representation of statistical significance is as follows: \**P* < 0.05; \*\**P* < 0.01; \*\*\**P* < 0.001; \*\*\*\**P* < 0.0001. Data showing statistical outcomes are represented as mean ± SEM. The above statistical analyses were performed using GraphPad Prism 6.0h. A Ridge Regression was performed for analyzing multiple linear regressions that suffer from co-linearity (plasma IL-10 and log_10_ HIV IntDNA) using P ≤ 0.20 to balance for type I and II errors.

All univariate group-differences were analyzed using a non-parametric Wilcoxon-ranked test. Whereas, all univariate correlation analyses were done using a non-parametric Spearman’s test. p<0.05 is reported as significant. MonteCarlo simulation approach implemented in the R package (https://cran.r-project.org/web/packages/MonteCarlo), was used to assess the significance of the overlap between *ex vivo* and *in vitro* pathways. Briefly, we simulated 1 million times 3 different lists of pathways having with the same length as the ex vivo and the in vitro pathways. Next, we assessed the number of times the overlap between these 3 lists is equal or higher than 47 using the MonteCarlo R function.

## Datasets Availability

The published article includes all datasets generated or analyzed during this study.

## Acknowledgements

We are grateful for the patients and clinical team.

## Author contributions

SPR and FPD conceptualized and conceived the experimental approaches. SPR, CNC, JH, CD, EM, DK, LL, XX, BT, JT and CS performed experiments in their area of expertise to address the major question. RB, VM, JPG, NK, CP, MP, SGD, JDE and RPS provided samples and supervised the execution of the major techniques on their labs, contributing to data plot and interpretation. LM, BH, DG, DH developed, synthetized and tested anti-IL-10 monoclonal antibody. MA, JPG provided the bioinformatic analysis and data integration. SPR and RPS wrote the manuscript. DHB, VM, CP, SGD, MP, JDE and RPS, contributed to critical review of the manuscript. All authors contributed to the manuscript development and have critically reviewed and approved the final version.

